# Active *E. coli* heteromeric acetyl-CoA carboxylase forms polymorphic helical tubular filaments

**DOI:** 10.1101/2024.05.28.596234

**Authors:** Xueyong Xu, Amanda Silva de Sousa, Griffin L. Martell, Trevor J. Boram, Sundharraman Subramanian, Wen Jiang, Jeremy R. Lohman

## Abstract

The *Escherichia coli* heteromeric acetyl-CoA carboxylase (ACC) has three functional subunits assumed to form an elusive catalytic complex and they are involved in allosteric and transcriptional regulation. The *E. coli* ACC represents almost all ACCs from pathogenic bacteria, making it a key antibiotic development target to fight growing antibiotic resistance. Furthermore, it is a model for cyanobacterial and plant plastid ACCs as biofuel engineering targets. Here, we report the catalytic *E. coli* ACC complex surprisingly forms tubes rather than dispersed particles. The cryo-EM structure reveals key protein-protein interactions underpinning efficient catalysis and how transcriptional regulatory roles are masked during catalysis. Discovering the protein-protein interaction interfaces that facilitate catalysis, allosteric, and transcriptional regulation provides new routes to engineering catalytic activity and new targets for drug discovery.

**One-Sentence Summary:** Bacterial heteromeric acetyl-CoA carboxylase forms tubes to promote efficient catalysis and mask transcriptional regulation.

## Main Text

As the first committed step in fatty acids biosynthesis, acetyl-CoA carboxylases (ACCs) are highly regulated. In the majority of prokaryotes, such as *Escherichia coli*, ACCs are heteromeric consisting of four proteins forming three functional subunits.^1^ End-products of fatty acid biosynthesis allosterically feedback regulate heteromeric ACCs, although the site of action is unknown.^2-4^ Some *E. coli* ACC subunits regulate or are proposed to regulate their own transcription,^5-7^ and one ACC subunit is known to interact with nitrogen assimilation regulatory protein GlnB,^8-10^ revealing broader transcriptional regulatory roles. However, the most important mode of prokaryotic heteromeric ACC regulation is through the formation of a distinct complex which is yet to be isolated and structurally characterized.^11^ Most plants have plastid-associated heteromeric ACCs, for which the *E. coli* ACC is a model system.^12^ The heteromeric *E. coli* ACC is necessary for growth and conserved in most bacterial pathogens, making them validated and promising antibiotic development targets.^13^ Thus, an *E. coli* ACC complex structure will provide knowledge essential for both antibiotic development and biofuel engineering.

Early studies reported that *E. coli*-like heteromeric ACC complexes readily disassociate upon cell lysis, which prevented experimentation on the native complex.^11,14-16^ Nevertheless, the three dissociated ACC subunits can be assayed individually or reconstituted into a complete system. In the first catalytic step, a 100 kDa homodimeric biotin carboxylase subunit (BC, AccC) uses ATP and bicarbonate to generate carboxy-biotin linked to a 17 kDa biotin-carboxy carrier protein (BCCP, AccB)(Fig. 1).^17-22^ In the second step, the carboxy-BCCP translocates to a 136 kDa heterotetrameric carboxyltransferase subunit (CT, AccA/AccD), where the carboxy group is transferred to acetyl-CoA to generate malonyl-CoA.^23^ In vitro studies demonstrated that protein:protein interactions (PPI) within the reconstituted ACC complex dramatically increase reaction kinetics and prevent the reverse reaction, establishing a regulatory role for complex formation.^16,24,25^ While crystal structures of the individual subunits are known, some disordered regions could not be modeled and are likely responsible for the PPIs (Fig. 1A).

**Fig. 1.**
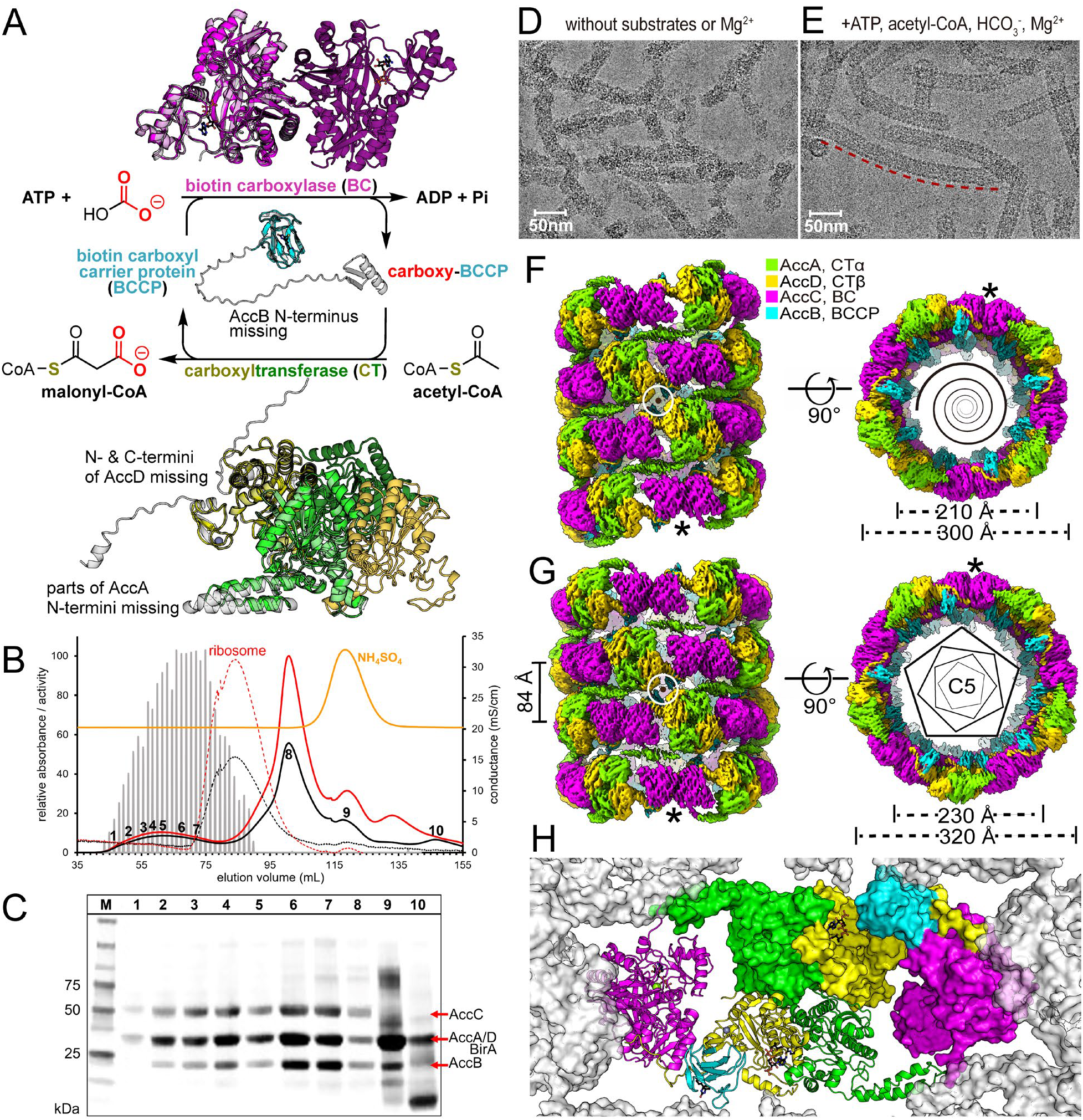
Catalytic activity, purification and cryo-EM structures of *E. coli* ACC. (**A**) Catalytic activity and structures of isolated *E. coli* ACC subunits. Two distinct catalytic activities of the ACC system are shown in the center. Previous crystal structures of AccC (PDB 1DV2) are colored magenta, AccB (PDB 1BDO) colored cyan and AccA:AccD system (PDB 2F9Y) colored green/yellow, respectively and their respective AlphaFold models (shown in light gray). Colors are carried through subsequent figures. The N-termini of AccA/B/D and C-termini of AccD are predicted in the AlphaFold models but missing in published crystal structures. The dimer complements of AccC and AccA:AccD are shown in a darker shade. (**B**) Size exclusion chromatography of the ACC complex and comparison to the *E. coli* ribosome resolved over a HiPrep 16/60 Sephacryl S-500 HR column (20 MDa mass cutoff). Absorbance at 280 nm (black) or 260 nm (red)for *E. coli* ACC expressed from pRSF as solid lines and *E. coli* ribosome (∼2.7 MDa) as thin dashed lines . Relative ACC activity is shown as gray bars for all fractions from 42 to 140 mL normalized to protein concentration. Conductance is shown as an orange line and indicates where ammonium sulfate from the ACC sample elutes, which indicates that some proteins have specific interactions with the column material slowing their elution, fraction 9 and 10.. (**C**) SDS-PAGE of fractions from SEC purification showing the presence of the ACC proteins. The numbers for the lanes correspond to numbered fractions in the chromatogram above. The amount of material in each lane is normalized to constant absorbance at 280 nm. (**D** and **E**) Representative micrographs of helical assemblies formed in the absence (D) and in the presence (E) of substrates and MgCl_2_, respectively. The tube highlighted by a red dashed line is longer than 0.3 μm. (**F** and **G**), Cryo-EM density maps of the narrow and the wide tubes, respectively. C5 point group symmetry of the wide tube is shown as regular pentagons and the axes of dihedral symmetries of both tubes are perpendicular to the helical axes and go through the centers of the white circles. Equivalent positions are indicated by ^*^. (**H**) Inside the tube looking out at one of the five sets of protomers making up a ring of (G). One protomer is represented as a surface and another as a ribbon.

Due to the importance of prokaryotic ACCs as the rate limiting step of fatty acid biosynthesis,^26^ we desired a picture of how the PPIs in the ACC complex regulates catalytic activity of the subunits. Here we report single-particle cryo-EM structures of active *E. coli* ACC, revealing the system surprisingly forms large polymorphic helical tubular filaments. Tube formation 1) ensures the BC and CT dimeric nature is satisfied in the complex, 2) places the active sites in close proximity, and 3) obscures parts of the subunits involved in transcriptional regulation.

### Production and catalytic activity of the complex

Based on reports that *E. coli* ACC complex purification from wild-type cells is notoriously difficult, we started with a high copy number pRSF overexpression plasmid containing an artificial *accA*/*accD*/*accB*/*accC*/*birA* operon, see Supplementary Materials for details and procedures. The artificial operon has a native ribosome binding site (RBS) between *accB*/*accC* and synthetic RBS between the other genes. There are discrepancies in the literature for the ratio of AccB and AccC produced from the native RBS. In one report, the native *accB*/*accC* RBS yields AccB at equimolar levels with AccC, AccA and AccD.^26^ In a subsequent report, in the absence of AccA and AccD, AccB is expressed at twice the level of AccC.^27^ Nevertheless, overexpressed ACC activity is readily detectable in crude lysate and absent without induction from our artificial operon. Classical ammonium sulfate precipitation yielded a fraction with improved activity (Fig. S1A and Table S1). Subsequent size exclusion chromatography (SEC) allowed purification of the ACC complex (Fig. S2). A simple BC:BCCP:CT dimeric complex is predicted to be 270 kDa and a previous report suggests a reconstituted complex is greater than 640 kDa which could be at least 4 of each subunit.^24^ Yet we isolate something much larger. Over an appropriate SEC column, ACC catalytic activity is spread between the 20 MDa exclusion limit down to the size of the *E. coli* ribosome (2,700 kDa, ∼20 nm diameter) and appears to be a 1:1:1:1 ratio of AccA:AccB:AccC:AccD (Fig. 1B/1C). This ratio is estimated using densitometry, in which we try to account for differences in staining efficiency and a propensity for full length AccB to form multiple bands in the absence of the complex (Fig. S3). During SEC excess AccB is separated from the complex, as the 30 % ammonium sulfate fraction has an AccB:AccC ratio of ∼2:1. The 30 % ammonium sulfate pellet has a specific activity of 0.93 µmol/min/mg enzyme (2.1 s^-1^) and the best SEC fraction (Fig. 1B fraction 5) has a specific activity of 5.0 µmol/min/mg enzyme (6.8 s^-1^) (Fig. S1C/S1D and Table S1).

Direct comparison with previously reconstituted systems is difficult as many have subunit ratios far from equal, or only monitor half-reactions.^16,28^ One study reports a specific activity of 5.6 µmol/min/mg enzyme based on 0.16 µg BC, but a limiting concentration of CT (0.32 µM BC, 0.24 µM CT, 2 µM BCCP).^24^ For comparison, yeast ACC has an activity of ∼2 µmol/min/mg (8.5 s^-1^) as a 500 kDa homodimer.^29^

The N-terminal 58 residues of AccA are partially ordered and N-terminal 24 residues of AccD are disordered in crystal structures, while the N-terminal region of AccB is also missing from structural studies.^23^ We deleted these regions to determine their role in complex formation and catalytic activity. In order to produce the mutant proteins, we had to change the expression vector to pCDF, a lower copy number vector to control toxicity, see supplemental text. We also altered the protein production and purification procedures to optimize the yield of activity for the complex from pCDF and also purified the complex from pRSF under the same conditions for comparison (Fig. S4). The complex behaves essentially identically from both plasmids when purified by SEC and the AccB subunit is completely biotinylated (Fig. S5 and S6 and Table S1). Deletion of *birA* from the pCDF construct lowers biotinylation to 70% but activity is essentially unchanged. Deletion of the first 58 residues of AccA from the pCDF construct yields a potential complex over SEC that is much smaller and loses about 90% of the specific activity. Deletion of the first 58 residues of AccB, or the first 23 or 52 residues of AccD leads to material with activity that is just above the detection limit in crude lysate (which could be due to complementation by the wild type chromosomal copy of the mutant subunit), but can’t be further purified. These mutants establish that the N-terminal extensions of AccA, AccB and AccD are necessary for full catalytic activity.

### Cryo-EM structure and overall atomic model

On cryo-EM grids in the absence of substrates, the *E. coli* ACC complex (Fig. S2 fractions 3-7, note the AccB for this fraction is estimated to be ∼60% biotinylated) formed tube-like assemblies that exhibited extremely poor order preventing structure determination (Fig. 1D). Upon incubation of the complex with ATP, acetyl-CoA, bicarbonate and Mg^2+^ for 3 minutes well-ordered tubes of various lengths were formed (Fig. 1E). To determine if the tube like-assemblies form in vivo, we collected cryo-electron tomography tilt-series of intact *E. coli* BL21(DE3) cells overexpressing ACC from the pRSF plasmid. This tilt series reveals long tubes that appear to run the length of the *E. coli* cells and are reminiscent of the active form of ACC from single particle analysis (compare Fig. 1E and Fig. S7 or Supplemental Movie 1). Cryo-EM movies of the well-ordered tubes provided two major types of 2D-classes with different diameters, ∼300 Å (narrow tube) or ∼320 Å (wide tube) (Fig. S8). Initial helical reconstruction of both tubes yielded 3D structures with overall resolutions of ∼4 Å (Fig. 1F/1G). See supplemental text and Figure S9 for helical indexing, Figure S10 for symmetry parameters, Figure S11 for example density, and Table S2 for detailed data collection and refinement statistics. The subunits could be unambiguously modeled into the density. The wide tube is essentially a set of stacked rings with 10 sets of BC:BCCP:CT subunits per ring with rings offset vertically by 84 Å and rotated -27.6° with respect to the central axis (Fig. 1G and S10C). The narrow tube is helical in nature, but can also be considered as layers of rings with 10 subunit sets packed with an overlap at the ends (Fig. 1F and S10D). In both forms, two sets of BC:BCCP:CT form a protomer, the core linear repeating unit (Fig 1H).

Functional relevance of the ACC tubes is supported by multiple observations. The tubes form in vivo under overexpression conditions, ruling out ammonium sulfate precipitation artifacts. There is a potential disorder-to-order transition of the complex upon the addition of substrates and Mg^2+^ but not Mg^2+^ alone (Fig. 1D and 1E). Efficient catalytic activity occurs in a complex that appears to be the size of one ring (5 protomers) or larger (Fig. 1B/1C), and mutants that disrupt ring formation and stacking are catalytically compromised. Finally, all the active sites are located on the inside of the tube making a reaction chamber that can potentially facilitate substrate channeling between active sites (Fig 1H and S12). The chamber is also near neighboring active sites in stacks above and below that can potentially contribute to efficient catalysis. While we find two tube types, they are likely functionally identical, as there are no significant structural differences between tube types when considering the local protomer environment and PPIs.

Local reconstruction yielded much improved maps for both narrow tubes (3.26 Å) and wide tubes (3.31 Å) which were used to build and refine the atomic models of ACC protomers (Fig. S13). There is modest density for CoA/acetyl-CoA in the CT and Mg^2+^·ADP in the BC active sites reflecting the fact our structure represents an average of the catalytic states (Fig S14). The BC has a subdomain known to have two conformations, open or closed over the ATP substrate. Cryo-EM density for the BC subdomain indicates it is in multiple conformations which are associated with multiple catalytic states (Fig. 2B and S15). Further detailed descriptions of the cryo-EM density and modeling of individual subunits can be found in the supplemental text, whereas we focus on the newly discovered PPIs below.

**Fig. 2.**
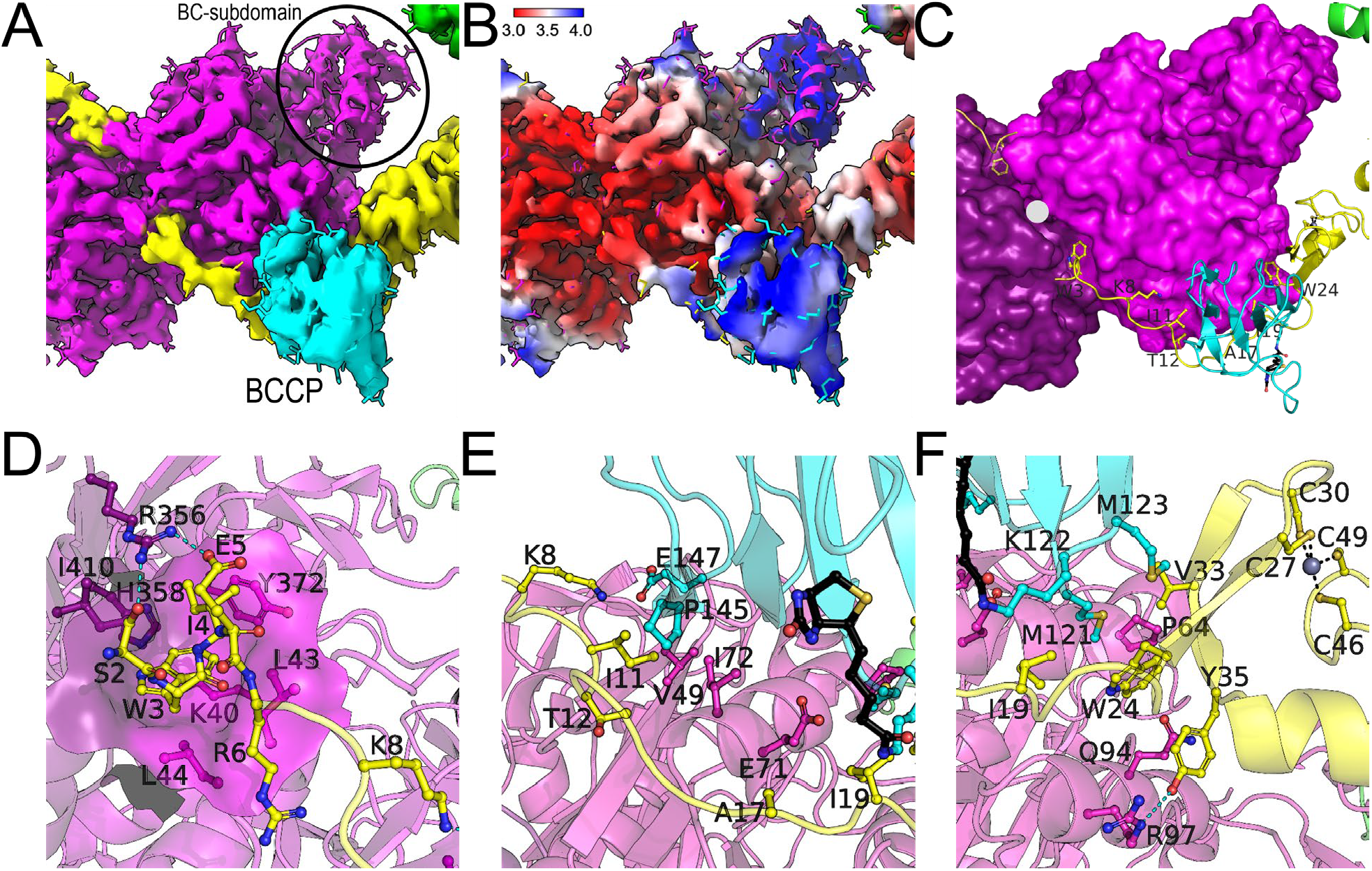
Detailed AccD interactions with BC and BCCP. (**A**) Density map of the helical tube colored according to the underlying model. (**B**) Density map of the helical tube colored by the local resolution. (**C**) Overview of the AccD interactions with BC and BCCP, the attached biotin is shown in black (note biotin density was weak and is modeled based on PDB 1BDO). Notice the two-fold symmetry at the BC interface with the two N-termini of AccD, indicated by a white ⬤. (**D**) The AccD Trp-3/Ile-4 binding pocket on the BC dimer interface. (**E**) Hydrophobic cluster near the middle of the extended AccD N-terminus. (**F**) Hydrophobic cluster between the AccD zinc-binding domain and BC/BCCP.

### ACC rings are generated by BC:AccD N-terminus interactions

Our structures reveal that the AccD N-terminal residues 2-23 and zinc-binding domain interacts with both the BC and BCCP (Fig. 2). Since the map was of high quality for the AccD N-terminus it reveals a network of van der Waals, hydrogen bonding and electrostatic interactions that stabilize the CT-BC-BCCP interaction. The AccD N-terminus interactions with BC and BCCP together generate 1582 Å^2^ of buried surface area (BSA) suggesting a stable complex. In comparison, the stable BC dimer interface is 1347 Å^2^ BSA. Even if the BCCP is displaced, the AccD:BC interactions alone generate 1145 Å^2^ BSA. The very N-terminus of AccD binds in a previously undocumented pocket on the BC dimeric interface (Fig. 2D). The importance of the AccD:AccC interactions at the BC dimer interface for promoting tube assembly may explain the previous observations that BC mutants deficient in dimerization fail to support *E. coli* growth at physiological concentration although the BC dimerization mutants are still catalytically active.^17,30,31^ In addition, we have revealed that deletion of the N-terminal 22 residues of AccD essentially abolishes activity (Fig. S4 and Table S1). This suggests the N-terminal AccD binding site on AccC can be a prime location for antibiotic design.

### Vertical interactions between rings via AccA N- and BC C-termini

The N-terminus of AccA consists of two alpha-helices (α1 residues 9-21 and α2 residues 37-57) that interact through regularly spaced hydrophobic residues in an antiparallel coiled-coils motif based on crystal structures and the AlphaFold model.^23,32^ Placing the AlphaFold model in our cryo-EM density reveals the N-terminal helices make close contacts across a pseudosymmetrical plane and only partially fit the density (Fig. S16). However, due to lower resolution around the AccA N-terminal helices it is difficult to unambiguously model the interactions. In some local 3D classifications, the N-terminal helices clearly interact in a domain-swapped manner, where α1 from one layer interacts with α2 from an adjacent tube layer (Fig. 3 and S17). Such interactions would act to create a stable tube, but may also play a role in catalytic regulation through long-range interactions with the CT active site. Deletion of the N-terminal helices decreases the catalytic activity of the complex ∼10-fold and abolishes formation of high mass complexes (Fig. S5), further establishing a role for these helices in stabilizing the active complex.

**Fig. 3.**
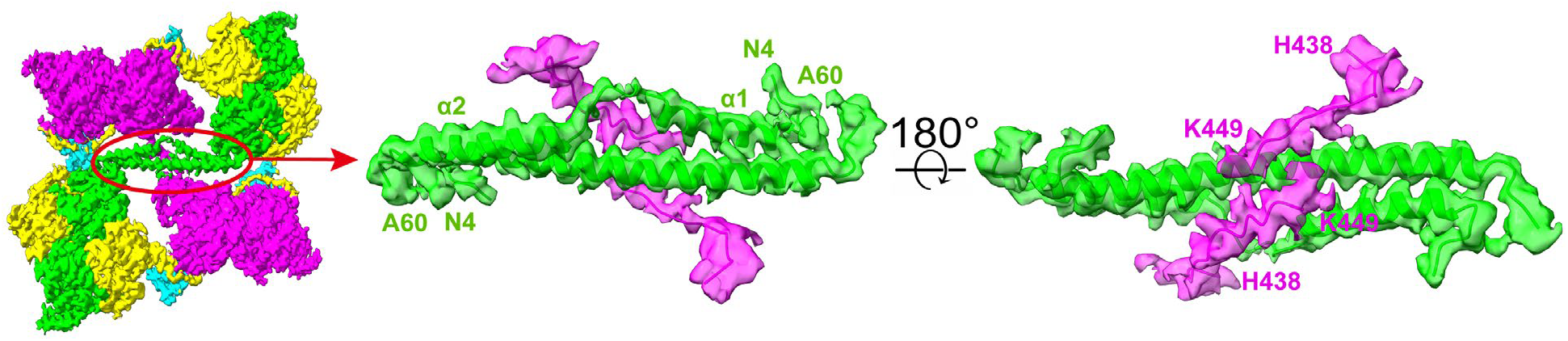
AccA N-terminus domain-swapped model and BC interactions. One of the possible sets of asymmetric interactions at the interface between stacks of rings. This figure illustrates how the tubes can be stabilized by novel interactions likely due to the amphipathic nature of the AccA N-terminal helices. AccA interactions with the BC domain C-terminus likely alter catalysis, since the BC C-terminal helix would need to locally unfold into an extended conformation.

The N-terminal α-helices of AccA interact with the BC C-terminus, likely further stabilizing the tubes. The BC C-terminus folds into a short helix (438-444) near the ATP binding site, with the terminal residues 445-449 in most crystal structures being disordered. Based on our cryo-EM maps, the C-terminus of the BC can be modeled in an extended conformation, placing hydrophobic residues Leu-444 and Leu-446 in contact with hydrophobic residues in the N-terminus of AccA (Fig. 3). Since the C-terminus of the BC is near the active site, the AccA:BC interaction can influence catalysis at the BC active site.

### BCCP translocation and avoidance of diffusion away from the tube

In the two-step ACC reaction, biotinylated BCCP must cycle between BC and CT, where the biotin moiety on BCCP is carboxylated by BC, then BCCP translocates to CT where the carboxyl group is transferred from carboxybiotin to acetyl-CoA (Fig. 1A, 1H and S12).^11^ There are plenty of gaps in the tube for small molecule substrates and products to flow in and out. At first glance, the biotinylated BCCP appears to be well situated inside the tube fixed to the wall at a non-catalytic site bound to the BC (previously seen in PDB 4HR7)(Fig. 4A) and AccD N-terminus (Fig. 3C/D).^33^ Together these interactions generate 869 Å^2^ BSA (AccD:BCCP 437 Å^2^, BC:BCCP 432 Å^2^), suggesting a moderately stable interaction. However, the BCCP is not at full occupancy (Fig. 2B), which is consistent with it in transit or bound to the catalytic active sites as our structure is a mixture of the various catalytic states. While in motion, the open ends of the tube provide a chance for the BCCP to diffuse away from the compartment. This raises the question, how is the BCCP retained in the tubes?

**Fig. 4.**
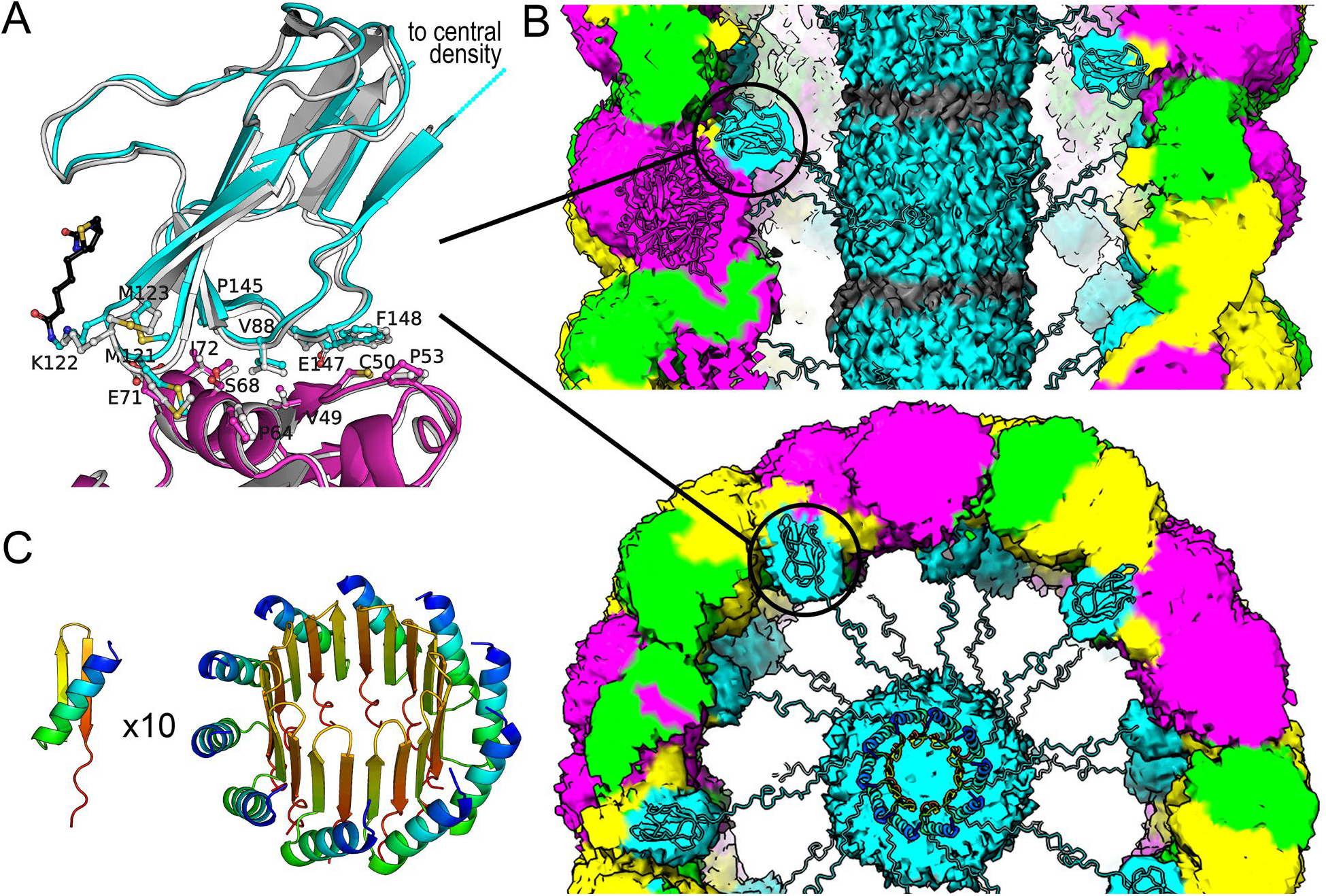
BCCP structure and interactions. (**A**) The interface between the BC and BCCP is shown for our cryo-EM structure, BCCP is in cyan with the attached biotin in black and BC in magenta, and the previous crystal structure in light gray. (**B**) The BCCP is near a central mass of density in our maps that can be explained by the BCCP N-terminus, which is modeled in the end on view. Linkers between the BCCP N-terminus and biotin-attachment domain are invisible in the cryo-EM maps but shown here for scale. (**C**) An AlphaFold3 model for 10 copies of the BCCP N-terminal 32 residues, which are predicted to fold into a β-barrel.

One answer is that the BC:BCCP interactions help to anchor the BCCPs, but that alone isn’t enough. The center of the tubes has cryo-EM density that likely corresponds to the BCCP N-terminus (Fig. 4B and S18). However, due to the diffuse nature of the central density atomic models can’t be built, so we rely solely on generative AI modeling in this region. AlphaFold3 predictions suggest that the first 30 N-terminal residues of the BCCP are folded into an α/β/β-motif (Fig. 4C).^34^ If 10 BCCP N-termini are modelled with AlphaFold3, the result is a β-barrel that generates ∼685 Å^2^ BSA at each interface, over 6000 Å^2^ total. Such a β-barrel corresponds well to the cryo-EM density in the center of the tubes (Fig. 4B). The diffuse cryo-EM density observed could be explained by the averaging of a mobile, off-center BCCP β-barrel structure (Fig. S19B). Other arrangements of the BCCP N-terminus could also account for the density in the center of the tube, such as two 10-membered β-barrels or a single 20-membered β-barrel (Fig. S19), see supplemental text for discussion of other models and caveats. Previous biophysical studies of the BCCP reveal it behaves as large aggregates alone and in the presence of the BC.^27,35,36^ The proposed BCCP N-terminal oligomeric β-barrel structure accounts for those previous cryptic observations.

Most importantly, this putative arrangement of the BCCP N-termini keeps the biotinylated C-terminal domains from freely diffusing away from the tubes. While some BCCP C-terminal domains are translocating, the other BCCPs hold them in position via interactions with their respective BC domains. In the single-chain dimeric eukaryotic ACCs, the BCCP is tethered to the other domains which abrogates the problem of diffusion of the BCCP away from each of the catalytic sites and creates a reaction chamber.^29,37^ The length and organization of the BCCP linkers in the single-chain homomeric system also creates allosteric interactions to regulate the overall catalytic activity. Here we demonstrate nature came up with a completely different architecture for ensuring the formation of a reaction chamber and BCCP anchoring mechanism in the heteromeric system.

### AccB N-terminus as a regulator of expression and activity in the tubular filament context

It is suggested that the BCCP of the *E. coli* ACC is expressed at roughly twice the number of the catalytic AccA/AccC/AccD proteins, based on multiple lines of evidence.^11^ Further evidence for double the amount of BCCP over BC and CT comes from one report, where the maximal activity of ACC required approximately twice as much BCCP as the catalytic domains.^25^ In our preparations from pRSF, the BCCP is expressed at double the BC. During SEC purification, excess BCCP is separated, suggesting it co-precipitates during ammonium sulfate fractionation rather than being bound in the complex (Fig S3). It may be expression of excess BCCP in wild-type *E. coli* is used to saturate the complex at equal ratios and excess BCCP is used in other regulatory pathways. One regulatory role of AccB is upregulation of biotin biosynthesis, where the presence of apo-AccB, alleviates biotin synthesis repression. Another regulatory role is sequestration of AccB by GlnB, linking fatty acid biosynthesis and nitrogen regulation.^9,10^ The N-terminus of AccB alone (residues 1-68) can negatively regulate expression of the *accB*/*accC* operon in *E. coli*, yet our structure suggests a biophysical role for the N-terminus.^5^ Early studies established in vitro *E. coli* ACC activity was dependent on high protein concentration.^38,39^ This suggests growth or cell-cycle dependent concentration changes of ACC in vivo can regulate activity. Measurements of *accB* mRNA and AccB protein levels of *E. coli* during growth indicate that as cell density increases and cells enter stationary phase, mRNA levels decrease but protein levels remain constant.^40^ The decrease in ATP and acetyl-CoA concentrations that accompanies the transition to stationary phase could drive an ACC order to disorder transition that we see in cryo-EM images in the absence of substrates (Fig. 1D and 1E). Such a transition could allow greater access to the BCCP N-terminus for transcriptional repression.

### Relevance of *E. coli* ACC structure to other heteromeric systems

A phylogenetic tree generated from concatenation of AccC/AccB/AccD/AccA from various organisms reveals two major clades (Fig. S20 and S21). The *E. coli* ACC is almost identical to species from pathogenic genus like *Brucella, Acinetobacter, Bordetella, Pseudomonas, Hemophilus, Vibrio, Yersinia, Salmonella, Klebsiella* and *Enterobacter*, with identities between 60-99%. Even cyanobacteria and plant plastidial ACCs are closely related to the *E. coli* ACC at greater than 49% and 45% identities, respectively. ACCs from *E. coli* and Gram-positive organisms share significant identity with an average around 45%. In some organisms, like *Acinetobacter baumannii* and *Lactobacillus amylolyticus*, the AccA N-terminal region is missing, suggesting their ACCs don’t form filaments, see supplementary text for details. This corroborates our observation that deletion of the *E. coli* AccA N-terminal region reduces but does not abolish activity. Conversely, the AccD N-termini and BC PPI residues essential for forming the *E. coli* ACC ring structures are highly conserved in other organisms.

### Higher-order polymer formation by ACCs

We have revealed that the heteromeric *E. coli* ACC should be added to the growing list of enzymes that form filaments.^41^ Human ACC contains all the domains on a single polypeptide, which forms a homodimer. The homodimer polymerizes into a filament in the presence of citrate or isocitrate to form an active complex. Another acyl-CoA carboxylase, 3-methylcrotonyl-CoA carboxylase from *Leishmania tarentola*, was also recently shown to form a filament in an inactive state.^42^ The observation of diverse acyl-CoA carboxylase oligomeric states suggests that filament formation is a broader feature of carboxyltransferase regulation across domains of life. Our demonstration that heteromeric complexes can be purified and structurally characterized, sets the stage to further examine the role of tubular filament formation and regulation for the heteromeric ACC family of enzymes.

## Conclusions

Our *E. coli* heteromeric ACC cryo-EM structures also raise many questions. For example, does the tube support crosstalk between the CT and BC active sites, which is suggested from kinetics experiments? Another question is, do *E. coli* ACC allosteric regulators like fatty acid bearing acyl carrier protein or GlnB, induce a disordered state as seen in the cryo-EM images without substrates? Experiments that allow examination of each tube catalytic state and dynamics are needed to provide that deeper level of understanding. It’s likely that cryo-EM tomography and the use of substrate analogs that prevent turnover will help achieve that deeper level of knowledge. Nevertheless, our cryo-EM structures provide a scaffold for modeling and testing allosteric mechanisms and address several lingering questions. The N-termini of AccA/B/D are functional rather than vestigial and play key roles in tube formation for setting up the catalytic active sites in close proximity. The AccD zinc-binding domain is buried in the complex making a putative role in translational repression unlikely and suggests the zinc-binding domain is primarily involved in PPIs.^6,7,24^ Bioinformatics along with our structures show the PPIs are likely conserved from prokaryotes to plants and set the stage to model plant ACC regulation for biofuel engineering. Finally, based on the uncovered PPIs unique to bacterial ACCs, our structures can be used to launch antibiotic drug discovery programs to combat the growing problem of antibiotic resistance.

## Acknowledgments

We would like to acknowledge the staff at the Michigan State University Mass Spectrometry and Metabolomics Core for assistance with data collection and processing. Some of this work was performed at the Midwest Center for Cryo-ET (MCCET) and the Cryo-EM Research Center located in the Department of Biochemistry at the University of Wisconsin-Madison, supported by the NIH Common Fund Transformative High Resolution Cryo-Electron Microscopy program (U24 GM139168). Work was also performed at the Michigan State University Cryo-EM Core Facility (RRID:SCR_028499)

## Author contributions

Conceptualization: JRL

Methodology: JRL, WJ, SS

Investigation: XX, AS, GM, TJB, SS

Visualization: XX, AS, GM, JRL

Funding acquisition: JRL, WJ

Project administration: JRL

Supervision: JRL, WJ

Writing – original draft: JRL, XX

Writing – review & editing: JRL, XX, AS, GM, TJB, SS, WJ

## Data and materials availability

The cryo-EM maps and coordinates for the *E. coli* ACC structures have been deposited in the Protein Data Bank and Electron Microscopy Data Bank. The depositions can be accessed with the following codes, PDB 9E4N/EMD 47516 for the narrow helical tube, PDB 9E4O/EMD 47517 for the wide ring stacked tube, PDB 8UZ2/EMD 42831 for the narrow helical tube protomer, and PDB 8UXZ/EMD 42790 for the wide ring stacked tube protomer. Models for the AccB N-termini and mass spectrometry data are deposited in Zenodo at https://doi.org/10.5281/zenodo.14743900 and 10.5281/zenodo.21267455, respectively.

Source data are provided with this paper.

## Supplementary Materials

Materials and Methods

Supplementary Text

Figs. S1 to S21

Table S1 and S2

