## Supplemental text and information for "Active *E. coli* heteromeric acetyl-CoA carboxylase forms polymorphic helical tubular filaments"

Materials and Methods

*Protein expression and purification*. An operon composed of *E. coli accA*/*accD*/*accB*/*accC*/*birA* with ribosome binding sites between each gene (native sequence between *accB*/*accC*) was synthesized and cloned into a modified pRSFDuet-1 vector by GenScript, see below for sequences (UniProt accession numbers: P0ABD5/P0A9Q5/P0ABD8/P24182/P06709). To ensure complete AccB biotinylation, *birA* was included in the operon. The plasmids were transformed into *E. coli* BL21(DE3) for protein production using electroporation and plated on LB with 50 µg/mL of kanamycin. A colony was picked and grown overnight in 50 ml LB supplemented with 50 µg/mL of kanamycin, 10 g/L glucose and 5 mM $MgCl_2$, and 500 µL of the culture was preserved as a 25% glycerol stock stored at -80°C. The expression strain was grown in 1 L of Terrific Broth with the addition of 50 µg/mL of kanamycin, 100 µM biotin and 0.2x trace metals (final concentrations: 10 µM $FeCl_3$, 4 µM $CaCl_2$, 2 µM $MnCl_2$, 2 µM $ZnSO_4$, 0.4 µM $CoCl_2$, 0.4 µM $CuCl_2$, 0.4 µM $NiCl_2$, 0.4 µM $Na_2MoO_4$, 0.4 µM $Na_2SeO_3$, 0.4 µM $H_3BO_3$). The cells were grown to an OD $A_{600}$ of ~1.2 at 37°C with shaking at 200 rpm, at which point the temperature was reduced to 18°C and upon thermal equilibrium, IPTG was added to 2 mM and shaking continued overnight. Cells were harvested by centrifugation at 5000 x g, then resuspended in buffer (100 mM Tris pH 8.0, 100 mM NaCl, 10% glycerol) and frozen. We used two purification methods, initially cells were thawed and ~ 1 mg/L of lysozyme and DNase were added followed by sonication on ice for 5 minutes with 10 s bursts and 30 s off intervals. The homogenized material was centrifuged at 4 °C for 20 minutes at 20,000 x g to pellet insoluble cell debris. The remaining supernatant was treated with solid ammonium sulfate (or a saturated ammonium sulfate solution containing 10% glycerol) to a concentration of 10% on ice and centrifuged again. The supernatant was treated with two further portions of ammonium sulfate to concentrations of 20 and 30%. Portions of the 30% ammonium sulfate pellets were either frozen for storage or resuspended in 100 mM Tris pH 8.0, 100 mM NaCl (resuspension buffer) and 500 µL of the resuspension at $A_{280}$ ~1.2 was applied to a Superose 6 10/300 GL size exclusion chromatography column (Cytiva) with fractionation ranges of ~5-5000 kDa and resolved using the same buffer at a flow rate of ~0.5 mL/min over an hour or analyzed for activity. In a second purification, to help reduce ribosome or RNA contamination, 1) we added 50 ng RNase A and 5 mM EDTA to the cell lysis procedure, 2) the ammonium sulfate fractionation was only done once at 30% and 3) the material was purified by SEC over the aforementioned Superose 6 column (which was used for cryo-EM analysis) or a HiPrep 16/60 Sephacryl S-500 HR column (Cytiva) with a fractionation range of ~20-0.03 MDa resolved with resuspension buffer at a flow rate of 0.25 mL/min over 6-8 hr. The *E. coli* 70S ribosome standard was purchased from NEB and thyroglobulin standard was from cytiva. The SEC fractions were examined for the presence of all the ACC subunits by SDS-PAGE with Flamingo (BioRad) staining, see below for conditions and quantification. Fractions containing the ACC complex were pooled and frozen in liquid nitrogen. The concentration of the complex was determined using an extinction coefficient of 73,780 $M^{-1}cm^{-1}$ at 280 nm based on a 1:1:1:1 complex of AccA:AccD:AccB:AccC. The concentration of the complex was also estimated using a Bradford or BCA assay with BSA as a calibration standard, which yielded similar values to the UV-Vis assay when fractions have reasonable 260/280 nm ratios..

The artificial operon was subcloned into a modified pCDFDuet-1 vector by GenScript and N-terminal deletion mutants were made from that construct and delivered as *E. coli* BL21(DE3) glycerol stocks. Due to trouble with production of some of the mutants we slightly altered the protocols to optimize expression from the pCDF plasmids and repeated the purification with pRSF as a control. The glycerol stocks were plated on fully defined non-inducing media plates (25 mM $Na_2HPO_4$, 25 mM $KH_2PO_4$, 50 mM $NH_4Cl$, 5 mM $Na_2SO_4$, 2 mM $MgSO_4$, 5.6 mM glucose, 7.5 mM aspartate, 0.2x trace metals [final concentrations 10 μM $FeCl_3$, 4 μM $CaCl_2$, 2 μM $MnCl_2$, 2 μM $ZnSO_4$, 0.4 μM $CoCl_2$, 0.4 μM $CuCl_2$, 0.4 μM $NiCl_2$, 0.4 μM $Na_2MoO_4$, 0.4 μM $Na_2SeO_3$, 0.4 μM $H_3BO_3$] and 200 mg/L of each amino acid except cysteine and tyrosine) with 50 μg/mL of streptomycin or kanamycin. A single colony was selected and inoculated into 50 mL of non-inducing minimal media (25 mM $Na_2HPO_4$, 25 mM $KH_2PO_4$, 50 mM $NH_4Cl$, 5 mM $Na_2SO_4$, 2 mM $MgSO_4$, 0.2x trace metals, 27.8 mM glucose and 18.8 mM aspartate) with 50 μg/mL of streptomycin or kanamycin and grown overnight. The growths were then inoculated and overexpressed in 1 L of Terrific Broth with 50 μg/mL of streptomycin or kanamycin. The cells were grown to an OD $A_{600}$ of ~1.8 at 37°C with shaking at 150 rpm. Once the desired OD was reached, the temperature and shaking were reduced to 18°C and 120 rpm, respectively. Upon thermal equilibrium, 0.4 mM IPTG, 400 μM biotin, 1 mM $MgSO_4$, and 1x trace metals were added to the culture, and shaking continued overnight. Cells were then harvested by centrifugation at 4000 x g, and the pelleted cells were resuspended in buffer (50 mM Tris pH 8.0, 50 mM NaCl, 10% glycerol). About 1 mg/L lysozyme and DNase were added to the cell suspension, and the mixture was sonicated on ice for 3 minutes with 10s on/off intervals. The homogenized material was centrifuged at 4 °C for 20 minutes at 20,000 x g to pellet insoluble cell debris. The supernatant was decanted and treated with a saturated ammonium sulfate solution to a concentration of 15% on ice then incubated for 15 minutes and centrifuged again at 20,000 x g. The ammonium sulfate precipitation step was repeated once to a final concentration of 40%. The ammonium sulfate pellets were resuspended in enough resuspension buffer (100 mM Tris pH 8.0, 100 mM NaCl) to fully solubilize. A 500 μl aliquot of the resuspended 40% ammonium sulfate fraction between an $A_{280}$ of 1 to 2 was applied to a Superose 6 Increase 10/300 GL size exclusion chromatography column (Cytiva) with a fractionation range of ~5-5000 kDa. The proteins were resolved using desalting buffer (10 mM Tris pH 8.0, 200 mM NaCl) at a flow rate of ~1.5 mL/min that adjusted to maintain a constant pressure. All ammonium sulfate and SEC fractions were quantified by a dual-wavelength Bradford assay with BSA as a calibration standard.[1] Samples were diluted to a similar total protein concentration for kinetic assay or in Laemmli loading buffer, heated to 90°C for 30 s then analyzed by SDS-PAGE using Any kD Mini-PROTEAN TGX Gels (BioRad), followed by Flamingo (BioRad) staining. The 260/280 ratios of fractions were highly variable due to nucleic acid contamination and/or the co-purification of adenosine nucleotides and CoAs.

The sequence of the pRSFDuet-1 or pCDFDuet-1 vectors carrying the *accA/D/B/C/birA* genes is shown below. The pRSF/pCDF sequence is shown in italics, the protein coding regions are underlined and colored as in Figure 1 with the *birA* coding region being uncolored. Artificial ribosome binding sites were included between *accA*/*accD*/, *accD*/*accB* and *accC*/*birA*, while the native *accB*/*accC* intergenic region and ribosome binding site was used (shown in bold). Regions corresponding to deletions have a strikethrough or double strikethrough. A similar operon in pCDF lacking *birA* was constructed by deleting the region between the *accC* and *birA* stop codons.

*TTTAATAaggagaTATACA*ATG~~AGTCTGAATTTCCTTGATTTTGAACAGCCGATTGCAGAGCTGGAAGCGAAAATCG~~
~~ATTCTCTGACTGCGGTTAGCCCGTCAGGATGAGAAACTGCATATTAACATCGATGAAGAAGTGCATCGTCTGCGTGAA~~

AAAAGCGTAGAACTGACACGTAAAATCTTCGCCGATCTCGGTGCATGGCAGATTGCGCAACTGGCACGCCATCCACA
GCGTCCTTATACCCTGGATTACGTTCGCCTGGCATTTGATGAATTTGACGAACTGGCTGGCGACCGCGCGTATGCAG
ACGATAAAGCTATCGTCGGTGGTATCGCCCGTCTCGATGGTCGTCCGGTGATGATCATTGGTCATCAAAAAGGTCGT
GAAACCAAAGAAAAAATTCGCCGTAACTTTGGTATGCCAGCGCCAGAAGGTTACCGCAAAGCACTGCGTCTGATGCA
AATGGCTGAACGCTTTAAGATGCCTATCATCACCTTTATCGACACCCCGGGGGCTTATCCTGGCGTGGGCGCAGAAG
AGCGTGGTCAGTCTGAAGCCATTGCACGCAACCTGCGTGAAATGTCTCGCCTCGGCGTACCGGTAGTTTGTACGGTT
ATCGGTGAAGGTGGTTCTGGCGGTGCGCTGGCGATTGGCGTGGGCGATAAAGTGAATATGCTGCAATACAGCACCTA
TTCCGTTATCTCGCCGGAAGGTTGTGCGTCCATTCTGTGGAAGAGCGCCGACAAAGCGCCGCTGGCGGCTGAAGCGA
TGGGTATCATTGCTCCGCGTCTGAAAGAACTGAAACTGATCGACTCCATCATCCCGGAACCACTGGGTGGTGCTCAC
CGTAACCCGGAAGCGATGGCGGCATCGTTGAAAGCGCAACTGCTGGCGGATCTGGCCGATCTCGACGTGTTAAGCAC
TGAAGATTTAAAAAATCGTCGTTATCAGCGCCTGATGAGCTACGGTTACGCGTAAAGaggagaATACTAGATGAGCT
GGATTGAACGAATTAAAAGCAACATTACTCCCACCCGCAAGGCGAGCATTCCTGAAGGGGTGTGGACTAAGTGTGAT
AGCTGCGGTCAGGTTTTATACCGCGCTGAGCTGGAACGTAATCTTGAGGTCTGTCCGAAGTGTGACCATCACATGCG
TATGACAGCGCGTAATCGCCTGCATAGCCTGTTAGATGAAGGAAGCCTTGTGGAGCTGGGTAGCGAGCTTGAGCCGA
AAGATGTGCTGAAGTTTCGTGACTCCAAGAAGTATAAAGACCGTCTGGCATCTGCGCAGAAAGAAACCGGCGAAAAA
GATGCGCTGGTGGTGATGAAAGGCACTCTGTATGGAATGCCGGTTGTCGCTGCGGCATTCGAGTTCGCCTTTATGGG
CGGTTCAATGGGGTCTGTTGTGGGTGCACGTTTCGTGCGTGCCGTTGAGCAGGCGCTGGAAGATAACTGCCCGCTGA
TCTGCTTCTCCGCCTCTGGTGGCGCACGTATGCAGGAAGCACTGATGTCGCTGATGCAGATGGCGAAAACCTCTGCG
GCACTGGCAAAAATGCAGGAGCGCGGCTTGCCGTACATCTCCGTGCTGACCGACCCGACGATGGGCGGTGTTTCTGC
AAGTTTCGCCATGCTGGGCGATCTCAACATCGCTGAACCGAAAGCGTTAATCGGCTTTGCCGGTCCGCGTGTTATCG
AACAGACCGTTCGCGAAAAACTGCCGCCTGGATTCCAGCGCAGTGAATTCCTGATCGAGAAAGGCGCGATCGACATG
ATCGTCCGTCGTCCGGAAATGCGCCTGAAACTGGCGAGCATTCTGGCGAAGTTGATGAATCTGCCAGCGCCGAATCC
TGAAGCGCCGCGTGAAGGCGTAGTGGTACCCCCGGTACCGGATCAGGAACCTGAGGCCTAAAGaggagaATACTAGA
TGGATATTCGTAAGATTAAAAAACTGATCGAGCTGGTTGAAGAATCAGGCATCTCCGAACTGGAAATTTCTGAAGGC
GAAGAGTCAGTACGCATTAGCCGTGCAGCTCCTGCCGCAAGTTTCCCTGTGATGCAACAAGCTTACGCTGCACCAAT
GATGCAGCAGCCAGCTCAATCTAACGCAGCCGCTCCGGCGACCGTTCCTTCCATGGAAGCGCCAGCAGCAGCGGAAA
TCAGTGGTCACATCGTACGTTCCCCGATGGTTGGTACTTTCTACCGCACCCCAAGCCCGGACGCAAAAGCGTTCATC
GAAGTGGGTCAGAAAGTCAACGTGGGCGATACCCTGTGCATCGTTGAAGCCATGAAAATGATGAACCAGATCGAAGC
GGACAAATCCGGTACCGTGAAAGCAATTCTGGTCGAAAGTGGACAACCGGTAGAATTTGACGAGCCGCTGGTCGTCA
TCGAG**TAAcgaggcgaacATG**CTGGATAAAATTGTTATTGCCAACCGCGGCGAGATTGCATTGCGTATTCTTCGTGC
CTGTAAAGAACTGGGCATCAAGACTGTCGCTGTGCACTCCAGCGCGGATCGCGATCTAAAACACGTATTACTGGCAG
ATGAAACGGTCTGTATTGGCCCTGCTCCGTCAGTAAAAAGTTATCTGAACATCCCGGCAATCATCAGCGCCGCTGAA
ATCACCGGCGCAGTAGCAATCCATCCGGGTTACGGCTTCCTCTCCGAGAACGCCAACTTTGCCGAGCAGGTTGAACG
CTCCGGCTTTATCTTCATTGGCCCGAAAGCAGAAACCATTCGCCTGATGGGCGACAAAGTATCCGCAATCGCGGCGA
TGAAAAAAGCGGGCGTCCCTTGCGTACCGGGTTCTGACGGCCCGCTGGGCGACGATATGGATAAAAACCGTGCCATT
GCTAAACGCATTGGTTATCCGGTGATTATCAAAGCCTCCGGCGGCGGCGGCGGTCGCGGTATGCGCGTAGTGCGCGG
CGACGCTGAACTGGCACAATCCATCTCCATGACCCGTGCGGAAGCGAAAGCTGCTTTCAGCAACGATATGGTTTACA
TGGAGAAATACCTGGAAAATCCTCGCCACGTCGAGATTCAGGTACTGGCTGACGGTCAGGGCAACGCTATCTATCTG
GCGGAACGTGACTGCTCCATGCAACGCCGCCACCAGAAAGTGGTCGAAGAAGCGCCAGCACCGGGCATTACCCCGGA
ACTGCGTCGCTACATCGGCGAACGTTGCGCTAAAGCGTGTGTTGATATCGGCTATCGCGGTGCAGGTACTTTCGAGT
TCCTGTTCGAAAACGGCGAGTTCTATTTCATCGAAATGAACACCCGTATTCAGGTAGAACACCCGGTTACAGAAATG
ATCACCGGCGTTGACCTGATCAAAGAACAGCTGCGTATCGCTGCCGGTCAACCGCTGTCGATCAAGCAAGAAGAAGT
TCACGTTCGCGGCCATGCGGTGGAATGTCGTATCAACGCCGAAGATCCGAACACCTTCCTGCCAAGTCCGGGCAAAA
TCACCCGTTTCCACGCACCTGGCGGTTTTGGCGTACGTTGGGAGTCTCATATCTACGCGGGCTACACCGTACCGCCG
TACTATGACTCAATGATCGGTAAGCTGATTTGCTACGGTGAAAACCGTGACGTGGCGATTGCCCGCATGAAGAATGC
GCTGCAGGAGCTGATCATCGACGGTATCAAAACCAACGTTGATCTGCAGATCCGCATCATGAATGACGAGAACTTCC
AGCATGGTGGCACTAACATCCACTATCTGGAGAAAAAACTCGGTCTTCAGGAAAAATAAAGaggagaATACTAGATG
AAGGATAACACCGTGCCACTGAAATTGATTGCCCTGTTAGCGAACGGTGAATTTCACTCTGGCGAGCAGTTGGGTGA
AACGCTGGGAATGAGCCGGGCGGCTATTAATAAACACATTCAGACACTGCGTGACTGGGGCGTTGATGTCTTTACCG
TTCCGGGTAAAGGATACAGCCTGCCTGAGCCTATCCAGTTACTTAATGCTAAACAGATATTGGGTCAGCTGGATGGC
GGTAGTGTAGCCGTGCTGCCAGTGATTGACTCCACGAATCAGTACCTTCTTGATCGTATCGGAGAGCTTAAATCGGG
CGATGCTTGCATTGCAGAATACCAGCAGGCTGGCCGTGGTCGCCGGGGTCGGAAATGGTTTTCGCCTTTTGGCGCAA
ACTTATATTTGTCGATGTTCTGGCGTCTGGAACAAGGCCCGGCGGCGGCGATTGGTTTAAGTCTGGTTATCGGTATC
GTGATGGCGGAAGTATTACGCAAGCTGGGTGCAGATAAAGTTCGTGTTAAATGGCCTAATGACCTCTATCTGCAGGA
TCGCAAGCTGGCAGGCATTCTGGTGGAGCTGACTGGCAAAACTGGCGATGCGGCGCAAATAGTCATTGGAGCCGGGA
TCAACATGGCAATGCGCCGTGTTGAAGAGAGTGTCGTTAATCAGGGGTGGATCACGCTGCAGGAAGCGGGGATCAAT
CTCGATCGTAATACGTTGGCGGCCATGCTAATACGTGAATTACGTGCTGCGTTGGAACTCTTCGAACAAGAAGGATT

~~GGCACCTTATCTGTCGCGCTGGGAAAAGCTGGATAATTTTATTAATCGCCCAGTGAAACTTATCATTGGTGATAAAG~~
~~AAATATTTGGCATTTCACGCGGAATAGACAAACAGGGGGCTTTATTACTTGAGCAGGATGGAATAATAAAACCCTGG~~
~~ATGGGCGGTGAAATATCCCTGCGTAGTGCAGAAAAATAA~~TCCGTAATCT*CCTCGAGTCTGGTAAAGA*

Excess untagged AccB was co-expressed from pCOLADuet-1 or pRSFDuet-1 with the *accA*/*D*/*B*/*C*/*birA* operon from pCDF to determine if AccB levels alter activity of the purified complex. The sequence of the pRSF/pCOLA vector carrying the *accB/birA* genes for coexpression with *accA*/*D*/*B*/*C*/*birA* genes in pCDF is shown below. The pRSF/pCOLA sequence is shown in italics, the protein coding regions are underlined.

*TTTAATAaggagaTATACA*ATGGATATTCGTAAGATTAAAAAACTGATCGAGCTGGTTGAAGAATCAGGCATCTCCG
AACTGGAAATTTCTGAAGGCGAAGAGTCAGTACGCATTAGCCGTGCAGCTCCTGCCGCAAGTTTCCCTGTGATGCAA
CAAGCTTACGCTGCACCAATGATGCAGCAGCCAGCTCAATCTAACGCAGCCGCTCCGGCGACCGTTCCTTCCATGGA
AGCGCCAGCAGCAGCGGAAATCAGTGGTCACATCGTACGTTCCCCGATGGTTGGTACTTTCTACCGCACCCCAAGCC
CGGACGCAAAAGCGTTCATCGAAGTGGGTCAGAAAGTCAACGTGGGCGATACCCTGTGCATCGTTGAAGCCATGAAA
ATGATGAACCAGATCGAAGCGGACAAATCCGGTACCGTGAAAGCAATTCTGGTCGAAAGTGGACAACCGGTAGAATT
TGACGAGCCGCTGGTCGTCATCGAGTAAAGaggagaATACTAGATGAAGGATAACACCGTGCCACTGAAATTGATTG
CCCTGTTAGCGAACGGTGAATTTCACTCTGGCGAGCAGTTGGGTGAAACGCTGGGAATGAGCCGGGCGGCTATTAAT
AAACACATTCAGACACTGCGTGACTGGGGCGTTGATGTCTTTACCGTTCCGGGTAAAGGATACAGCCTGCCTGAGCC
TATCCAGTTACTTAATGCTAAACAGATATTGGGTCAGCTGGATGGCGGTAGTGTAGCCGTGCTGCCAGTGATTGACT
CCACGAATCAGTACCTTCTTGATCGTATCGGAGAGCTTAAATCGGGCGATGCTTGCATTGCAGAATACCAGCAGGCT
GGCCGTGGTCGCCGGGGTCGGAAATGGTTTTCGCCTTTTGGCGCAAACTTATATTTGTCGATGTTCTGGCGTCTGGA
ACAAGGCCCGGCGGCGGCGATTGGTTTAAGTCTGGTTATCGGTATCGTGATGGCGGAAGTATTACGCAAGCTGGGTG
CAGATAAAGTTCGTGTTAAATGGCCTAATGACCTCTATCTGCAGGATCGCAAGCTGGCAGGCATTCTGGTGGAGCTG
ACTGGCAAAACTGGCGATGCGGCGCAAATAGTCATTGGAGCCGGGATCAACATGGCAATGCGCCGTGTTGAAGAGAG
TGTCGTTAATCAGGGGTGGATCACGCTGCAGGAAGCGGGGATCAATCTCGATCGTAATACGTTGGCGGCCATGCTAA
TACGTGAATTACGTGCTGCGTTGGAACTCTTCGAACAAGAAGGATTGGCACCTTATCTGTCGCGCTGGGAAAAGCTG
GATAATTTTATTAATCGCCCAGTGAAACTTATCATTGGTGATAAAGAAATATTTGGCATTTCACGCGGAATAGACAA
ACAGGGGGCTTTATTACTTGAGCAGGATGGAATAATAAAACCCTGGATGGGCGGTGAAATATCCCTGCGTAGTGCAG
AAAAATAATCCGTAATCT*CCTCGAGTCTGGTAAAGA*

For in vitro addition of holo-AccB to the ACC complex, AccB was expressed from a modified pRSF vector that co-expressed *accB* and *birA* in an operon, with the N-terminus of AccB carrying a His-tag and TEV-protease site for His-tag removal. Expression of AccB was carried out as above, but the cell pellet was resuspended in resuspension buffer containing 20 mM imidazole. After sonication and centrifugation, the crude tagged AccB was loaded on a HisTrap FF 5 mL column (Cytiva), washed with resuspension buffer with 20 mM imidazole and lacking glycerol, followed by elution with the same buffer containing 250 mM imidazole. Fractions containing tagged-AccB were desalted into anion exchange buffer containing 10 mM Tris pH 8.0, 10 mM NaCl over a HiPrep 26/10 desalting column (Cytiva). The desalted material was either treated with TEV protease overnight before further purification or immediately loaded onto a HiPrep Q HP 16/10 anion exchange chromatography column (Cytiva). Proteins were eluted with a gradient from 10 mM Tris pH 8.0 with 10 mM NaCl to 500 mM NaCl over 10 column volumes. His-tagged holo-AccB eluted between 400-500 mM NaCl, which was concentrated, frozen as pellets in liquid nitrogen and stored at -80°C. A large portion of the fractions containing AccB co-purified with BirA. Furthermore, freeze-thaw cycles of purified AccB leads to a change in migration on SDS-PAGE with smaller bands appearing, so subsequent experiments for AccB in-gel quantification were carried out on freshly purified protein. We altered the protocol by adding 6 M urea to the wash buffer for HisTrap purification which effectively removed BirA. To polish the His-tagged sample, the eluted holo-AccB was desalted into 50 mM Tris pH 8.0, 50 mM NaCl then immediately loaded onto the anion exchange chromatography column and eluted with a gradient from 50 mM Tris pH 8.0 with 50 mM NaCl to 500 mM NaCl over 10 column volumes. His-tagged holo-AccB eluted between 300-400 mM NaCl. Although holo-AccB contains no tryptophan it does have two tyrosines so UV/Vis could be used to calculate concentrations (extinction coefficient of 2980 $M^{-1}cm^{-1}$ at 280 nm). When the his-tag is not cleaved the intact protein has an extra tyrosine increasing the extinction coefficient (4470 $M^{-1}cm^{-1}$ at 280 nm). The presence of biotin on AccB was confirmed by intact protein LC/MS.

The sequence of the modified pRSF vector carrying the *accB/birA* genes for expression and purification of holo-AccB is shown below. The pRSF-Duet1 sequence is shown in italics, the protein coding regions are underlined, and a TEV protease recognition coding sequence is in bold. When TEV protease is used to remove the N-terminal His-6 tag from the overexpressed protein it leaves an N-terminal serine at position 0.

*TTTAATAaggagaTATACCatgggcagcagccatcaccatcatcaccacagc***ggatccgagaacctctacttccaaA**<br>**GT**ATGGATATTCGTAAGATTAAAAAACTGATCGAGCTGGTTGAAGAATCAGGCATCTCCGAACTGGAAATTTCTGAA<br>GGCGAAGAGTCAGTACGCATTAGCCGTGCAGCTCCTGCCGCAAGTTTCCCTGTGATGCAACAAGCTTACGCTGCACC<br>AATGATGCAGCAGCCAGCTCAATCTAACGCAGCCGCTCCGGCGACCGTTCCTTCCATGGAAGCGCCAGCAGCAGCGG<br>AAATCAGTGGTCACATCGTACGTTCCCCGATGGTTGGTACTTTCTACCGCACCCCAAGCCCGGACGCAAAAGCGTTC<br>ATCGAAGTGGGTCAGAAAGTCAACGTGGGCGATACCCTGTGCATCGTTGAAGCCATGAAAATGATGAACCAGATCGA<br>AGCGGACAAATCCGGTACCGTGAAAGCAATTCTGGTCGAAAGTGGACAACCGGTAGAATTTGACGAGCCGCTGGTCG<br>TCATCGAGTAAAGaggagaATACTAGATGAAGGATAACACCGTGCCACTGAAATTGATTGCCCTGTTAGCGAACGGT<br>GAATTTCACTCTGGCGAGCAGTTGGGTGAAACGCTGGGAATGAGCCGGGCGGCTATTAATAAACACATTCAGACACT<br>GCGTGACTGGGGCGTTGATGTCTTTACCGTTCCGGGTAAAGGATACAGCCTGCCTGAGCCTATCCAGTTACTTAATG<br>CTAAACAGATATTGGGTCAGCTGGATGGCGGTAGTGTAGCCGTGCTGCCAGTGATTGACTCCACGAATCAGTACCTT<br>CTTGATCGTATCGGAGAGCTTAAATCGGGCGATGCTTGCATTGCAGAATACCAGCAGGCTGGCCGTGGTCGCCGGGG<br>TCGGAAATGGTTTTCGCCTTTTGGCGCAAACTTATATTTGTCGATGTTCTGGCGTCTGGAACAAGGCCCGGCGGCGG<br>CGATTGGTTTAAGTCTGGTTATCGGTATCGTGATGGCGGAAGTATTACGCAAGCTGGGTGCAGATAAAGTTCGTGTT<br>AAATGGCCTAATGACCTCTATCTGCAGGATCGCAAGCTGGCAGGCATTCTGGTGGAGCTGACTGGCAAAACTGGCGA<br>TGCGGCGCAAATAGTCATTGGAGCCGGGATCAACATGGCAATGCGCCGTGTTGAAGAGAGTGTCGTTAATCAGGGGT<br>GGATCACGCTGCAGGAAGCGGGGATCAATCTCGATCGTAATACGTTGGCGGCCATGCTAATACGTGAATTACGTGCT<br>GCGTTGGAACTCTTCGAACAAGAAGGATTGGCACCTTATCTGTCGCGCTGGGAAAAGCTGGATAATTTATTAATCG<br>CCCAGTGAAACTTATCATTGGTGATAAAGAAATATTTGGCATTTCACGCGGAATAGACAAACAGGGGGCTTTATTAC<br>TTGAGCAGGATGGAATAATAAAACCCTGGATGGGCGGTGAAATATCCCTGCGTAGTGCAGAAAAATAATCCGTAATC<br>T*CCTCGAGTCTGGTAAAGA*

Similar to holo-AccB, AccC was expressed from a modified pRSF vector with the N-terminus of AccC carrying a His-tag and TEV-protease site for His-tag removal. Expression of AccC was carried out as above, but without the addition of biotin. The concentration of AccC was determined using UV-Vis with an extinction coefficient of 27,850 $M^{-1}cm^{-1}$ at 280 nm.

The sequence of the pRSF vector carrying the *accC* gene for expression and purification is shown below. The pRSF-Duet1 sequence is shown in italics, the protein coding regions are underlined, and a TEV protease recognition coding sequence is in bold. TEV protease was used to remove the N-terminal $His_6$ tag from the overexpressed protein leaving an N-terminal SG before Met-1.

*TTTAATAaggagaTATACCatg*ggcagcagccatcaccatcatcaccacagc**ggatccgagaacctctacttccaaA**<br>**GC**GGGATGCTGGATAAAATTGTTATTGCCAACCGCGGCGAGATTGCATTGCGTATTCTTCGTGCCTGTAAAGAACTG<br>GGCATCAAGACTGTCGCTGTGCACTCCAGCGCGGATCGCGATCTAAAACACGTATTACTGGCAGATGAAACGGTCTG<br>TATTGGCCCTGCTCCGTCAGTAAAAAGTTATCTGAACATCCCGGCAATCATCAGCGCCGCTGAAATCACCGGCGCAG

TAGCAATCCATCCGGGTTACGGCTTCCTCTCCGAGAACGCCAACTTTGCCGAGCAGGTTGAACGCTCCGGCTTTATC
TTCATTGGCCCGAAAGCAGAAACCATTCGCCTGATGGGCGACAAAGTATCCGCAATCGCGGCGATGAAAAAAGCGGG
CGTCCCTTGCGTACCGGGTTCTGACGGCCCGCTGGGCGACGATATGGATAAAAACCGTGCCATTGCTAAACGCATTG
GTTATCCGGTGATTATCAAAGCCTCCGGCGGCGGCGGCGGTCGCGGTATGCGCGTAGTGCGCGGCGACGCTGAACTG
GCACAATCCATCTCCATGACCCGTGCGGAAGCGAAAGCTGCTTTCAGCAACGATATGGTTTACATGGAGAAATACCT
GGAAAATCCTCGCCACGTCGAGATTCAGGTACTGGCTGACGGTCAGGGCAACGCTATCTATCTGGCGGAACGTGACT
GCTCCATGCAACGCCGCCACCAGAAAGTGGTCGAAGAAGCGCCAGCACCGGGCATTACCCCGGAACTGCGTCGCTAC
ATCGGCGAACGTTGCGCTAAAGCGTGTGTTGATATCGGCTATCGCGGTGCAGGTACTTTCGAGTTCCTGTTCGAAAA
CGGCGAGTTCTATTTCATCGAAATGAACACCCGTATTCAGGTAGAACACCCGGTTACAGAAATGATCACCGGCGTTG
ACCTGATCAAAGAACAGCTGCGTATCGCTGCCGGTCAACCGCTGTCGATCAAGCAAGAAGAAGTTCACGTTCGCGGC
CATGCGGTGGAATGTCGTATCAACGCCGAAGATCCGAACACCTTCCTGCCAAGTCCGGGCAAAATCACCCGTTTCCA
CGCACCTGGCGGTTTTGGCGTACGTTGGGAGTCTCATATCTACGCGGGCTACACCGTACCGCCGTACTATGACTCAA
TGATCGGTAAGCTGATTTGCTACGGTGAAAACCGTGACGTGGCGATTGCCCGCATGAAGAATGCGCTGCAGGAGCTG
ATCATCGACGGTATCAAAACCAACGTTGATCTGCAGATCCGCATCATGAATGACGAGAACTTCCAGCATGGTGGCAC
TAACATCCACTATCTGGAGAAAAAACTCGGTCTTCAGGAAAAATAATCCGTAATTCT*CCTCGAGTCTGGTAAAGA*

The AccA/AccD subunit was expressed from a modified pRSF vector with the N-terminus of AccA carrying a His-tag and TEV-protease site for His-tag removal. Expression of AccA/D was carried out as above, but again without the addition of biotin. The concentration of AccA/D was determined using UV-Vis with an extinction coefficient of 42,860 $M^{-1}cm^{-1}$ at 280 nm.

The sequence of the pRSF vector carrying the *accA/accD* genes for expression and purification is shown below. The pRSF-Duet1 sequence is shown in italics, the protein coding regions are underlined, and a TEV protease recognition coding sequence is in bold. TEV protease was used to remove the N-terminal $His_6$ tag from the overexpressed protein leaving an AccA N-terminus beginning with Ser2.

*TTTAATAaggagaTATACCatg*ggcagcagccatcaccatcatcaccacagc**ggatccgagaacctctacttccaa**A
GTCTGAATTTCCTTGATTTTGAACAGCCGATTGCAGAGCTGGAAGCGAAAATCGATTCTCTGACTGCGGTTAGCCGT
CAGGATGAGAAACTGGATATTAACATCGATGAAGAAGTGCATCGTCTGCGTGAAAAAAGCGTAGAACTGACACGTAA
AATCTTCGCCGATCTCGGTGCATGGCAGATTGCGCAACTGGCACGCCATCCACAGCGTCCTTATACCCTGGATTACG
TTCGCCTGGCATTTGATGAATTTGACGAACTGGCTGGCGACCGCGCGTATGCAGACGATAAAGCTATCGTCGGTGGT
ATCGCCCGTCTCGATGGTCGTCCGGTGATGATCATTGGTCATCAAAAAGGTCGTGAAACCAAAGAAAAAATTCGCCG
TAACTTTGGTATGCCAGCGCCAGAAGGTTACCGCAAAGCACTGCGTCTGATGCAAATGGCTGAACGCTTTAAGATGC
CTATCATCACCTTTATCGACACCCCGGGGGCTTATCCTGGCGTGGGCGCAGAAGAGCGTGGTCAGTCTGAAGCCATT
GCACGCAACCTGCGTGAAATGTCTCGCCTCGGCGTACCGGTAGTTTGTACGGTTATCGGTGAAGGTGGTTCTGGCGG
TGCGCTGGCGATTGGCGTGGGCGATAAAGTGAATATGCTGCAATACAGCACCTATTCCGTTATCTCGCCGGAAGGTT
GTGCGTCCATTCTGTGGAAGAGCGCCGACAAAGCGCCGCTGGCGGCTGAAGCGATGGGTATCATTGCTCCGCGTCTG
AAAGAACTGAAACTGATCGACTCCATCATCCCGGAACCACTGGGTGGTGCTCACCGTAACCCGGAAGCGATGGCGGC
ATCGTTGAAAGCGCAACTGCTGGCGGATCTGGCCGATCTCGACGTGTTAAGCACTGAAGATTTAAAAAATCGTCGTT
ATCAGCGCCTGATGAGCTACGGTTACGCGTAAAGaggagaATACTAGATGAGCTGGATTGAACGAATTAAAAGCAAC
ATTACTCCCACCCGCAAGGCGAGCATTCCTGAAGGGGTGTGGACTAAGTGTGATAGCTGCGGTCAGGTTTTATACCG
CGCTGAGCTGGAACGTAATCTTGAGGTCTGTCCGAAGTGTGACCATCACATGCGTATGACAGCGCGTAATCGCCTGC
ATAGCCTGTTAGATGAAGGAAGCCTTGTGGAGCTGGGTAGCGAGCTTGAGCCGAAAGATGTGCTGAAGTTTCGTGAC
TCCAAGAAGTATAAAGACCGTCTGGCATCTGCGCAGAAAGAAACCGGCGAAAAAGATGCGCTGGTGGTGATGAAAGG
CACTCTGTATGGAATGCCGGTTGTCGCTGCGGCATTCGAGTTCGCCTTTATGGGCGGTTCAATGGGGTCTGTTGTGG
GTGCACGTTTCGTGCGTGCCGTTGAGCAGGCGCTGGAAGATAACTGCCCGCTGATCTGCTTCTCCGCCTCTGGTGGC
GCACGTATGCAGGAAGCACTGATGTCGCTGATGCAGATGGCGAAAACCTCTGCGGCACTGGCAAAAATGCAGGAGCG
CGGCTTGCCGTACATCTCCGTGCTGACCGACCCGACGATGGGCGGTGTTTCTGCAAGTTTCGCCATGCTGGGCGATC
TCAACATCGCTGAACCGAAAGCGTTAATCGGCTTTGCCGGTCCGCGTGTTATCGAACAGACCGTTCGCGAAAAACTG
CCGCCTGGATTCCAGCGCAGTGAATTCCTGATCGAGAAAGGCGCGATCGACATGATCGTCCGTCGTCCGGAAATGCG
CCTGAAACTGGCGAGCATTCTGGCGAAGTTGATGAATCTGCCAGCGCCGAATCCTGAAGCGCCGCGTGAAGGCGTAG
TGGTACCCCCGGTACCGGATCAGGAACCTGAGGCCTAATCCGTAATTCT*CCTCGAGTCTGGTAAAGA*

The individual subunits AccA/AccD, AccB and AccC were applied to a Superose 6 Increase 10/300 GL size exclusion chromatography column and resolved using resuspension buffer (100 mM Tris pH 8.0, 100 mM NaCl).

*SDS-PAGE analysis and densitometry.* Gels for routine analysis were either Mini-PROTEAN TGX 4-10% or Any kD precast gels (Bio-Rad). To concentrate some samples trichloroacetic acid precipitation and acetone washes were performed. To determine the response of proteins to Flamingo stain, gels were imaged using a ChemiDoc MP (BioRad) imaging system and analyzed with Image Lab 6.0.1 to determine volumes. The AccB and AccC proteins had the same linear response to the flamingo dye over the range tested from 0.37-10 µg. In general, AccB had the best response to the flamingo dye with overnight (~16 hr) fixing, 2-4 hour staining and no or very short destaining. For gels not used in densitometry, images were captured by the ChemiDoc MP, a GelDoc Go (BioRad) with a Blue Tray or Typhoon (GE) imager, the imaging parameters were not optimized to avoid overexposure with these instruments, so the images are only qualitative.

*Intact protein mass spectrometry.* The intact ACC proteins were analyzed by LC/MS using a Waters Xevo G2-XS QToF interfaced with a Waters Acquity UPLC. The samples (10 µL at ~10 µM) were injected onto a Waters Acquity UPLC protein BEH C4 column (2.1 x 100 mm, 300Å, 1.7 µm) and after flushing with 0.1 formic acid in water with 2% acetonitrile for 3 min, the proteins were eluted using a gradient to 30% acetonitrile over 1 min followed by a gradient to 55% acetonitrile over 5 minutes with subsequent steps to wash the column by increasing the acetonitrile to 75% over a minute then holding for 2 minutes and finally holding at 2% acetonitrile for 3 minutes. The column temperature was 40°C and the flow rate was 0.3 ml/min. Proteins were ionized by electrospray operating in positive ion mode with capillary voltage at 3 kV, cone voltage at 35 V, source temp 100°C, desolvation temp at 350°C, desolvation gas flow was 600 L/hr and cone gas flow was 25 L/hr. Mass spectra were acquired in continuum mode with a 1 second scan time across an m/z range of 200-2000. Protein mass spectra were deconvoluted to give a neutral mass of the intact proteins using Masslynx software and the MaxEnt I algorithm in the range of 14000-55000 Da and 15 cycles of refinement which lead to convergence. Two water blank injections were run between samples to minimize carryover.

*Assay for ACC activity*. Typical assays contained 250 µM acetyl-CoA (depending on supplier, this material had 5-10% CoA contamination), 4 mM ATP, 50 mM sodium bicarbonate and between 2-200 µg of ACC complex from various purification steps in a buffer of 10 mM $MgCl_2$, 100 mM bis-tris propane pH 7.0, 20 mM KCl. The reactions were quenched with formic acid to approximately pH 2 at various time points over 60 min. For relative activity, a single timepoint was used where there was approximately 50% conversion for the most active sample, and all fractions tested contained the same amount of ACC complex based on 280 nm absorbance. For specific activity, multiple timepoints and/or ACC complex concentrations were used to find a point where there was ~10% conversion of acetyl-CoA to malonyl-CoA. HPLC over a Luna 5 µm C18(2) 100 Å 250 × 4.6 mm column (Phenomenex) was used to monitor the conversion of acetyl-CoA to malonyl-CoA with a gradient of water with 0.5% TFA to 20% acetonitrile over 20 minutes at a flow rate of 1 mL/min. Under these conditions CoA elutes at ~ 9 min, malonyl-CoA at ~ 10.5 min and acetyl-CoA at 11 min. Peak areas monitored at 254 nm were integrated to determine % conversion. For crude lysate, acetyl-CoA hydrolysis by metabolic enzymes competes with ACC carboxylation, such that wild-type cells appear to be lacking activity. ACC overexpression merely provides enough ACC to outcompete the background hydrolysis rates.

*Single particle cryo-EM grid preparation and data collection*. Initially the ACC complex purified over the Superose 6 10/300 GL column (Fig. S2 fractions 3-7) was diluted to 1.5 mg/ml with 50 mM HEPES pH 7.5, 150 mM NaCl and 3 μL of the sample was applied to the gold side of a glow-discharged Quantifoil R1.2/1.3, 400 mesh grid coated with a thin layer of gold. The grid was subsequently blotted for 14 s at 4°C, 100% relative humidity and plunge frozen into liquid ethane using a Vitrobot Mark IV system. Sample grids with better ordered tubes were subsequently obtained by using 2.5 mg/ml ACC complex in 50 mM HEPES pH 7.5, 100 mM bicarbonate, 7.5 mM ATP, 20 mM $MgCl_2$ and 1 mM acetyl-CoA for 3 min at room temperature before plunge-freezing into liquid ethane with the same grid and procedure as the former sample.

High-throughput data collection of the well-ordered helical tubes was performed on a Titan Krios G4 300 kV transmission electron microscope equipped with a post-column Gatan Quantum GIF energy filter and a Gatan K3 direct electron detector. Movie stacks were recorded automatically using the EPU software at a nominal magnification of 105K in super-resolution mode with a pixel size of 0.411 Å per super-resolution pixel at the specimen level. The zero-loss slit width of the energy filter was set to 20 eV. A total dose of 54.438 e/Å$^2$ was fractionated into 50 frames with a defocus range set between -0.7 μm and -2 μm.

*Single particle cryo-EM image processing*. Image processing was performed with the CryoSPARC software,[2] except that RELION 4.0 was used for 3D classification,[3] and HI3D was used to determine helical symmetry parameters.[4] Super-resolution movies were 2× Fourier binned, gain-normalized, dose-weighted, aligned and corrected for beam-induced motion using Patch Motion Correction. CTF was estimated using Patch CTF Estimation. The tubes were first manually picked with a subset of micrographs, then the templates generated by 2D classification were used to automatically pick the tubes in all micrographs using Filament Tracer. The contaminant picks and highly curved tubes were removed by Inspect Picks. The helical segments were extracted, and 2 rounds of 2D classification were performed to clean up the segments. Then the segments which showed 2 different diameters in 2D classes were selected to reconstruct 3D structures separately using the same following steps. The initial models were built using asymmetric Helical Refinement in which helical symmetry was not applied. Helical symmetry parameters were determined from the asymmetric reconstructions with HI3D in real space. From the 2D lattices generated by the auto-correlation function (ACF) of the cylindrical projections of the initial models of the tubes in HI3D, helical twists (x-coordinate) and rises (y-coordinate) were determined by the unit cell vectors with the shortest distances to the equators (Fig. S9A/B). Then the particles in cryoSPARC cs format files were converted to RELION-compatible star files using images2star.py in the JSPR software [5] and were input into RELION to perform 3D classification with helical parameter search. Particles in classes that yielded reconstructions at high resolution with nearly identical helical symmetry were selected and imported back into CryoSPARC. The 3D helical refinement was performed with local helical parameter search. In addition, for each asymmetric unit, both $(BC)_2(BCCP)_2$ and $(CT\alpha, CT\beta)_2$ have a C2 symmetry perpendicular to the helical axis, which adds an additional dihedral point group symmetry to both narrow and wide tubes. Therefore, additional D1 symmetry of the narrow tube and D5 symmetry of the wide tube were imposed during helical refinement. The resolution of the final reconstruction was estimated based on the gold-standard Fourier shell correlation (FSC) = 0.143 criterion between independently refined half-sets split at the filament level. To calculate a local reconstruction without imposing helical symmetry, new particle datasets were created by symmetry expansion from helical symmetry and dihedral symmetry, i.e., duplicating the original particles' orientation and position parameters for each asymmetry unit in the helical lattice. The sub-volume corresponding to one protomer, $(BC)_2(BCCP)_2(CT\alpha, CT\beta)_2$, was selected to generate masks. Local 3D classifications were performed, and the best classes were selected for local 3D refinement. The masked local reconstructions of the asymmetric units had nominal resolutions of 3.26 Å for the narrow tube and 3.31 Å for the wide tube, respectively. To increase the high-resolution details in the maps, all the maps including the whole tubes and local protomer maps were sharpened with the B factors reported for the reconstructions. To figure out if the helix swapped interaction of N-terminal α-helices of CTα (residues 4-57) exists, focused 3D classification was performed without alignment using the same masks as that used in the local reconstructions.

*Atomic modeling*. The models of narrow and wide tubes were built based on the 3.26 Å and 3.31 Å maps from the local reconstructions. The crystal structures of CT tetramer (PDB ID: 2F9Y), BCCP (PDB ID: 1BDO), and BC in complex with Mg-ADP and bicarbonate (PDB ID: 3RV4) were used as starting models to rigid-body fit into the density maps using Chimera.[6] Then the models were manually extended and adjusted in Coot.[7] Final models were refined in Phenix using real-space refinement with appropriate restraints.[8] Buried surface area was determined using PISA.[9]

*Cryo-electron tomography*. The E. coli BL21(DE3) with ACC operon in pRSF was grown in 5 mL of LB with 50 μg/mL of kanamycin and biotin until an $OD_{600}$ of about 0.6 at which point 2 mM IPTG was added and growth continued for 30 minutes. The bacteria were spun down and resuspended to a final $OD_{600}$ of 0.2 in PBS buffer containing 10 nm gold fiducials coated with BSA, applied to R 3.5/1 Quantifoil grids, and then plunge frozen using a Vitrobot Mark IV (blot time: 3.0 seconds and blot force: -2). Tilt series were collected at the Midwest Center for Cryo-ET (MCCET) on a Titan Krios G3i equipped with a K3 direct electron detector and operating at 300 keV with a post-column GIF (20 eV slit width). Tilt series images were recorded between ± 51° with dose-symmetric image collection every 3° at 53,000 X nominal magnification (2.138 Å/pixel). Images were collected for 0.138 s (8 frames/tilt image), resulting in 2.65 electrons/$Å^2$ per tilt image (90 electrons/$Å^2$, total exposure dose). Individual micrographs were corrected for motion using MotionCor2,[10] and the corrected images were stitched back into a tilt series using the newstack functionality in IMOD.[11] Tilt series alignment using fiducial markers was carried out using IMOD.[11]

Supplementary Text:

**Complex production**
We made attempts to increase the activity of our isolated complex. We attempted co-expression of the ACC operon from pCDF (~20-40 copy number) with extra AccB being supplied by pRSF (~200 copy number). However, only a few co-transformant colonies were obtained, and the cell pellets were very small after overnight induction, and ACC activity wasn't readily detectable. This indicates that excess AccB is toxic when highly overexpressed in the presence of moderate

ACC operon expression. An imbalance of ACC subunit ratio has been reported to be toxic or cause plasmid instability before.[12] Co-transformation of the ACC operon lacking *birA* in pCDF with *accB*/*birA* in pCOLA (~20-40 copy number) yielded many co-transformants. These cells produced the ACC complex, but the activity in crude lysate was no better than pCDF carrying the ACC operon alone.

We initially attempted to produce complexes with AccA and AccD N-terminal deletion mutants from pRSF. However, transformations of the mutants in pRSF yielded few colonies in *E. coli* BL21(DE3) even with glucose supplementation for catabolite repression. The colonies that grew were small in size and produced very small pellets in subsequent overexpression experiments. SDS-PAGE of the overexpression cell lysate did not reveal overexpressed ACC subunits. Switching the ACC mutants to a pCDF plasmid along with catabolite repression or the use of non-inducing minimal media for plates and overnight starter cultures yielded relatively normal growth and allowed overexpression.

**Mass spectrometry analysis**

We analyzed SEC fractions I to IV (Fig. S5) and purified AccB by intact protein mass spectrometry over a C4 column to help partially separate subunits for better analysis by MaxEnt deconvolution. For his-tagged AccB we expect the N-terminal methionine to be removed giving a mass of 18854 Da (1%) and with biotinylation a mass of 18985 Da, based on MaxEnt analysis 99% is biotinylated (Fig. S6A). Upon treatment of the his-tagged holo-AccB with TEV protease, we get the expected mass of 17001 Da for the protein having a serine before the initiating methionine. Freeze-thawing the AccB sample a single time leads to the formation of smaller bands by SDS-PAGE. However, this treatment does not lead to readily detectable fragments by intact protein mass spectrometry, suggesting there are conformational changes that convert the aberrantly large band (~20 kDa) for AccB to one with a more appropriate size ~15 kDa considering it has an excess negative charge.

In certain cases, fraction I behaved poorly, so our analyses focused on fractions II-IV. For fractions II and III we could readily detect all subunits, except for the AccA N-terminal Δ58 mutant. For AccA we expect and found the N-terminal methionine to be removed, giving a mass of 35110 Da. For AccD we expect and found the N-terminal methionine to be removed, giving a mass of 33190. For AccC we expect and found the N-terminal methionine to be retained, giving a mass of 49321 Da. For AccB we expect and found retention of the N-terminal methionine giving a mass of 16687 Da for apo and 16913 Da for holo. In general, the ratio of apo to holo-AccB remained constant across fractions II-IV, and we summarize the results of the relative abundance for apo to holo-AccB in Supplemental Table S1. We would like to note that the results should not be viewed as quantitative since converting apo-AccB to holo-AccB converts a positively charged lysine to a neutral charge, which likely decreases instrument sensitivity as data were collected in positive mode, leading to an underestimation of biotinylation.

**Helical parameter and symmetry descriptions of the tubes**

The particles assigned to the two types of tubes were selected to perform helical 3D reconstruction. The structures show overall resolutions of 4.04 Å for the narrow and 3.98 Å for the wide tubes. The helical symmetry parameters of rise/twist are 19.2 Å/81.1° for the narrow tube and 84.2 Å/-27.6° for the wide tubes (Fig. S9). The narrow tube displays an additional D1 symmetry and the wide tube D5 symmetry with the dihedral axis perpendicular to the helical axes. These two types of tubes, though with drastically different helical rise/twist parameters and point group symmetries, are polymorphs of helical tubes formed from the same underlying 2D lattice of the ACC proteins but using different patches of the lattice to roll up into the tubes (Fig. S10), as often observed for helical structures.[13]

**Structure of the CT subunit**

Previously reported crystal structures served as initial models and were docked as rigid bodies into the cryo-EM maps. The 3.2 Å crystal structure of the *E. coli CT* $(AccA/AccD)_2$ heterotetramer was placed unambiguously into the maps,[14] along with an AlphaFold CT heterotetramer model.[15] The well-resolved EM densities and the AlphaFold model provided confidence for making some minor and major adjustments to the crystal structure. In the previous *E. coli* CT crystal structure, residues 4-57 of AccA had extremely high B-factors and were only partially built. Based on the cryo-EM maps, we were able to build the N-terminal region of AccA in various conformations (Fig. S16 and S17).

We found well-resolved density for residues 2-23 of AccD, which were disordered in the previous crystal structure. These residues form extensive PPIs. Mutations that disrupt the zinc-finger domain abolish CT activity without disrupting CT heterotetramer formation. [16,17] In the previous *E. coli* AccD crystal structure residues 93-98 were unmodeled and residues 86-93 were modeled with high B-factors.[14] Based on our well-resolved density and AlphaFold model, we could model these residues with high confidence and our model for residues 86-98 diverge significantly from the previous crystal structure. The 86-93 residues are near the acetyl-CoA binding site giving them a likely role in catalysis. The C-terminal residues 285-304 of AccD could not be resolved, leaving their function unclear, but they point toward the BC implicating a potential role in regulation.

Density for CoA is clearly identifiable, with some weak density for the acetyl-group of acetyl-CoA (Fig. S14A). Our structures support that the acetyl-CoA binding pocket is in the AccA/AccD heterodimeric interface as previously suggested.[14] Acetyl-CoA inserts deeply into the pocket and forms a network of interactions with the residues from both AccA and AccD around the pocket (Fig. S14A). The density for reactive acetyl-thioester is somewhat ambiguous and the thioester oxygen can be modeled as interacting with AccD Ala165 NH or AccD Gly205 NH or AccA Ser205 OH. The lack of density for the acetyl-group suggests the ligand isn't tightly bound in a productive conformation until carboxy-biotin BCCP binds and generates a ternary complex. There is no indication that the BCCP is bound here in our maps.

**Structure of the BC-BCCP subunits**

There are 27 *E. coli* BC domain crystal structures in the PDB that can fit our cryo-EM maps. Those crystal structures depict the BC domain in an open (PDB 1BNC, 1DV1)[18,19] or closed conformation (PDBs 1DV2, 2J9G, 2VR1, 3G8C, 3G8D, 3RUP, 3RV3, 3RV4) [20-22] where a small "B-domain" (residues 131-204) covers the ATP/bicarbonate binding site. The best fit structures are in the closed conformation, with PDB 3RV4 (ATP bound to R16E mutant) having the most similar small "B-domain" conformation (Fig. S15).[22] The "B-domain" density is weaker than the rest of the BC (Fig. 2B), reflecting multiple conformations due to the complex undergoing catalysis during grid preparation. Accordingly, there is density for a partially occupied

$Mg^{2+}$·ADP product in the BC active site (Fig. S14B). In the deposited local reconstruction, the BC C-terminus was only modeled to residue Leu-446 in a conformation similar to crystal structures. However, the entire BC C-terminus (His-438 to Lys-449) could be modeled in alternative conformations at lower occupancy but interacting with AccA (Fig. 3).
A previous structure of a BC-BCCP heterotetramer (PDB 4HR7)[23] revealed that the BC had a non-catalytic BCCP binding site. The same BC-BCCP interactions are found in our structure (Fig. 2C and 4A). The cryo-EM density for the BCCP is much weaker than the adjacent BC, but similar to the density for the BC "B-domain" (Fig. 2C and S13C/H). The weaker density suggests the BCCP is not fully occupied, which may be due to partial occupancy at the BC or CT active sites. While our SDS-PAGE analysis reveals AccB is full length, only the biotinylated C-terminal domain (residues 77-156) can be unambiguously modeled.[24] The density for the biotin prosthetic group and Lys-122 side chain are relatively weak, partly due to incomplete biotinylation, so we modeled the biotin orientation based on a high-resolution crystal structure (PDB 1BDO).[24]

**Detailed AccD N-terminal interactions with BC and BCCP**
The AccD residues Trp-3/Ile-4 form hydrophobic interactions with a hydrophobic pocket made of Lys-40/Leu-43/Leu-44/Tyr-372 of one BC subunit and His-358/Ile-410 from the adjacent BC subunit (Fig 2D). The AccD hydroxyl of Ser-2 and carboxylate of Glu-5 interacts with the guanidinium of Arg-356 (Fig 2D). The AccD N-terminal residues Lys-8 to Tyr-35 and the zinc-finger domain interact with both the BC and BCCP (Fig. 2C-2D). A salt bridge is formed between AccD Lys-8 and BCCP Glu-147. The rest of the interactions are hydrophobic in nature. A hydrophobic cluster is formed between AccD residues Ile-11/Thr-12, BC residues Val-49/Ile-172, and BCCP Pro-145 (Fig 2E). A second hydrophobic cluster is formed between AccD residues Ala-17/Ile-19/Trp-24/Val-33, BC residues Glu-71/Pro-64, and BCCP residues Met-121/Met-123 (Fig 2F). The AccD zinc-finger domain residues Trp-24/Tyr-35 interact with BC residues Gln-94/Arg-97. Mutation of the AccD zinc-finger cysteines to alanine abolish catalytic activity, likely through disruption of the AccD:BC interface. [16,17]

**Structure and role of the BCCP N-termini**
AlphaFold 2 and 3 both predict an αββ motif for the 35 N-terminal residues of the BCCP (average Cα pLDDT confidence score is 79). When multiple copies of the BCCP N-termini are input into AlphaFold 3, different results are seen depending on copy number. With 7 copies or less all models are of extended linear β-sheets. With 8 copies only one of the 5 models output is a β-barrel with ~630 $Å^2$ BSA between the monomers, while the other four models are linear β-sheets with ~677 $Å^2$ BSA between monomers. With 9 copies, 4 of the output models generate a β-barrel with ~640 $Å^2$ BSA between the monomers, while one output model is a linear β-sheet with ~685 $Å^2$ BSA. With 10 copies or more (up to 20 copies) all models are β-barrels. The 10 copy β-barrel has monomer interfaces with ~685 $Å^2$ BSA, while the 20 copy β-barrel has monomer interfaces with ~640 $Å^2$ BSA. Taken together these models suggest that ~10 subunits generate a β-barrel with fully buried monomer interfaces. When purified to homogeneity, the full length BCCP can form large aggregates, while the C-terminal domain is monomeric.[25,26] The aggregates can be directly attributed to the N-terminal 87 residues and specifically to the first 32 residues.[27] An AccB/AccC subcomplex can be overexpressed and purified, except when any four consecutive residues are deleted between residue 1 and 32 of AccB.[27] However, deleting regions of the linker between the predicted AccB αββ motif and C-terminal domain are tolerated,

suggesting this region is at least partially flexible.[28] In our modelling, the linker between the BCCP N-terminal 35-residues and C-terminal carrier domain can extend up to 150 Å if stretched out linearly, over half of the inner radius of the tubes, 230 Å (Fig S19A/B). Our model explains why deletions in the AccB linker regions are tolerated.[28]

**Conservation of PPI residues**

The residues at the interaction interfaces represented by AccD:BC are largely conserved (Fig. S21). The N-terminus of AccC Trp-3 and Ile-4 are mostly conserved in the Gram-negative bacteria and are large bulky hydrophobic residues, Leu/Phe, in Gram-positive bacteria. Those bulky residues are surrounded by mostly hydrophobic residues Leu-43, Leu-44, Tyr 372, Ile-410 and have van der Waals interactions with Lys-40 and His-358. Those hydrophobic pocket residues of the BC are again conserved in Gram-negative, whereas Lys-40/His-358 are replaced by hydrophobic residues in Gram-positive bacteria. The conservation at this pocket suggests it plays a common role in the regulation of BC activity. The main anchor of the AccD zinc-finger:BC interaction is AccD Trp-24, which is almost completely conserved. The residues IPEGV leading up to Trp-24 are relatively conserved in the β/γ-proteobacteria, but differ for the Gram-positives with significant similarities. The other AccD zinc-finger residues interacting with the BC Val-33 and Tyr-35 are also almost completely conserved. The BC residues interacting with the AccD zinc-finger include two helices, Ile-63, Pro-64, Ile-67, Glu-71, and Gln-94, Arg-97, Ser-98, Phe-100, which are very conserved among the β/γ-proteobacteria. Some of the Gram-positive bacteria exchange Glu-71 for a hydrophobic residue, while some Gram-positive and negatives trade Gln-94 for a hydrophobic residue, Arg-97 and Ser-98 are only conserved in the β/γ-proteobacteria, while Phe-100 is always a large hydrophobic residue. This reveals that the interactions with residues 63-71, which are closer to the BC:BCCP interface are more conserved, indicative of higher conservation for a BC:BCCP docking site in the complex. The BCCP primarily interacts with the AccD:BC interface with residues Met-121 and Met-123, which flank the bioinylated Lys-122, which are all completely conserved. A minor interaction between the three subunits is mediated by the BCCP Pro-145 and Glu-147, both of which are almost completely conserved. The BC:BCCP interface is further moderately conserved with respect to the BCCP hydrophobic contacts Val-88 and Phe148 with BC Val-49/Ile-51/Ala-65/Ile-72 and Pro-53, respectively. The hydrophobic contacts and moderate conservation in the BC:BCCP interface suggest drugs could be designed to target this region to destabilize the complex in an organism specific fashion.

The N-termini of AccA are primarily responsible for stabilizing the tubes through interaction between layers of rings or helical stacks. These N-termini are missing from a few genera, namely *Lactobacillus*, *Acinetobacter*, *Enterococcus* and *Streptococcus*. Nevertheless, in the majority of genera the N-terminal helices of AccA are present and have some conserved features, residues 5-22 are almost completely conserved, along with residues 34-61 having a common pattern of hydrophobic and hydrophilic residues. The lack of the AccA N-terminal helices from some species suggesting formation of distinct single ring assemblies and novel modes of allosteric regulation.

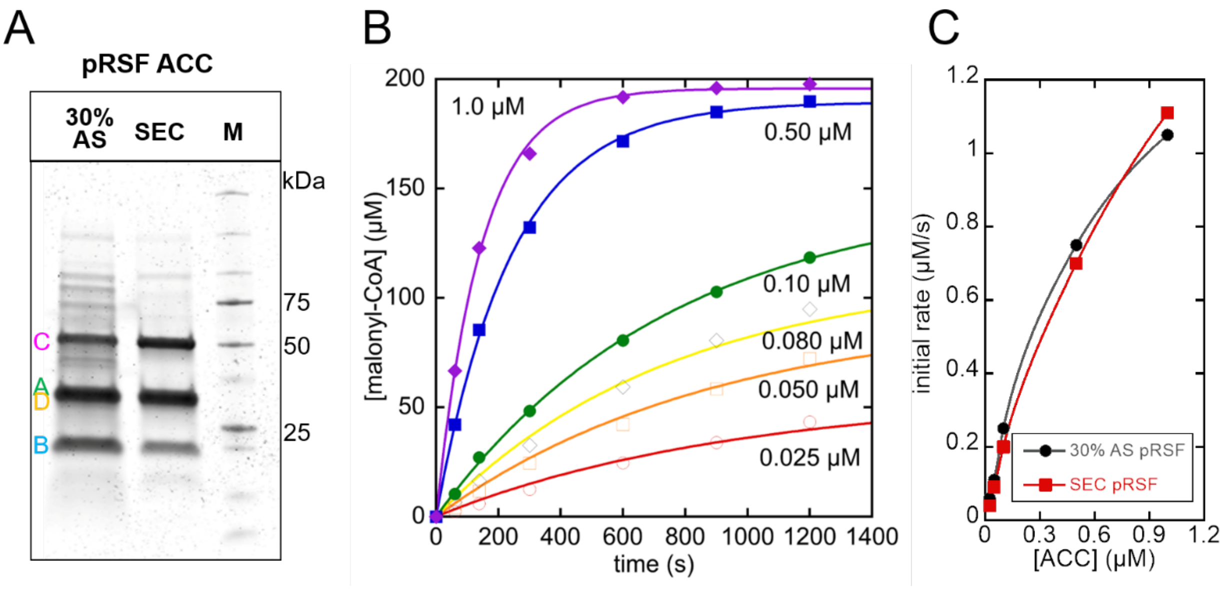


Figure S1. Catalytic activity and purity of ACC. (**A**) Comparison of purity for the 30% AS fraction and SEC fraction. SDS-PAGE gel stained with Flamingo. (**B**) Example reaction progress curves as a function of ACC concentration for a pooled SEC fraction, determined by HPLC analysis. These data were fit using a single exponential decay equation. In this experiment, the initial acetyl-CoA concentration was ~ 210 μM and CoA was ~40 μM due to contamination in the commercial material. (**C**) Initial rate vs ACC concentration from 30% AS pellet or SEC (for this experiment, fractions were pooled leading to about 4-fold lower activity than the best fraction). Trendlines are not fits to an equation and are only to indicate general trends. This experiment reveals an almost linear dependence on ACC complex concentration from pRSF. The slight curvature at higher ACC concentrations is due to a small number of data points during the steady state phase leading to a larger underdetermination of the initial rate, see panel B.

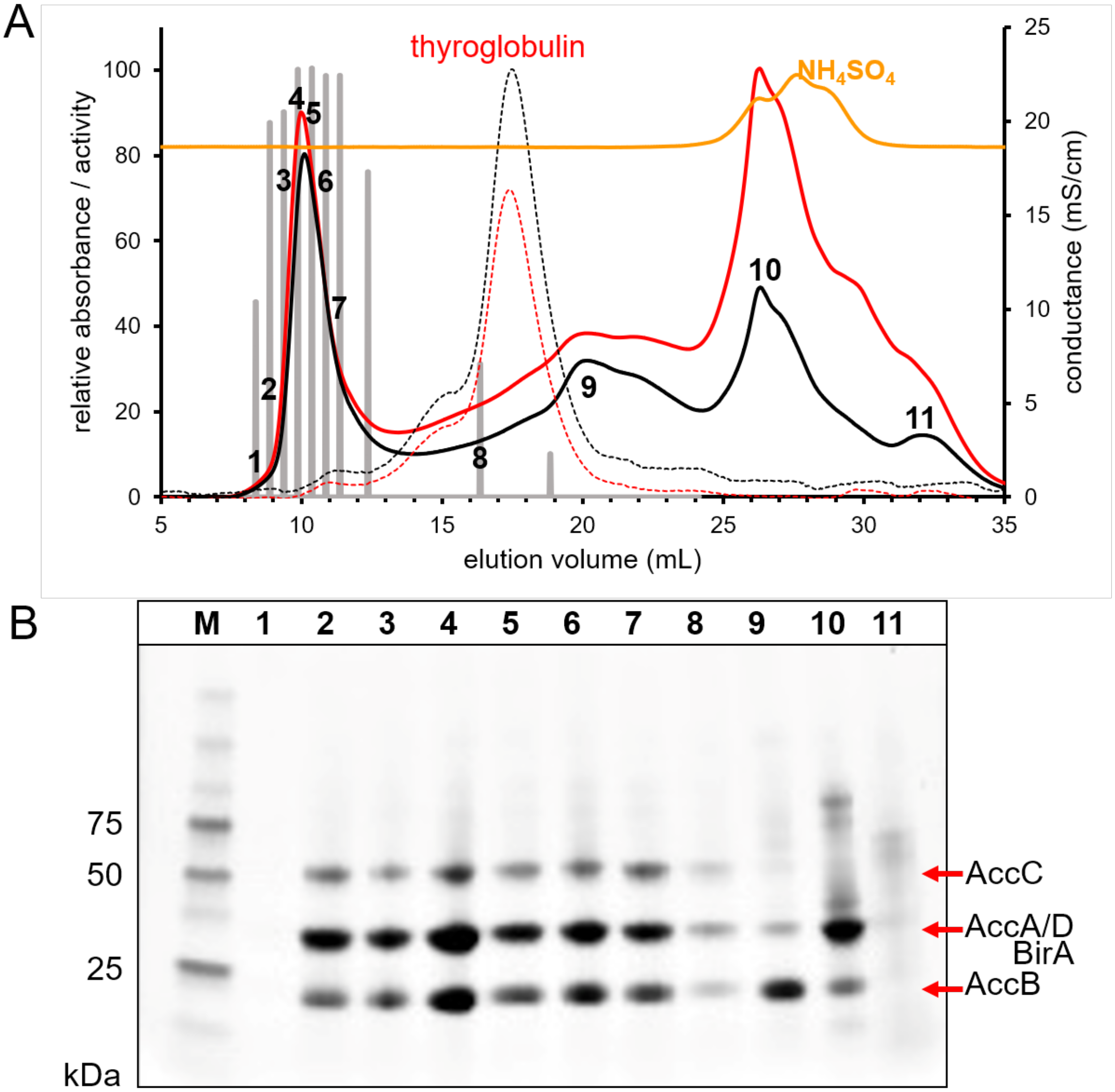


Figure S2. *E. coli* ACC size exclusion chromatography purification, activity and purity of the fractions from the pRSF plasmid. (**A**) Superose 6 10/300 GL column (5 MDa mass cutoff) purification result. Relative activity is shown in gray bar format for selected fractions normalized to 280 nm absorbance. The black lines are 280 nm relative absorbance, red lines are 254 nm relative absorbance and orange line is conductance. The solid black/red solid lines are for the ACC sample and thin dashed lines arefor a thyroglobulin standard (660 kDa). The ACC complex elutes just behind the void volume. In some fractions there is a significant contribution from nucleic acid absorbance as judged by a high 260/280 ratio. (**B**) SDS-PAGE of fractions from Superose 6 purification showing the presence of the ACC proteins. The numbers for the lanes correspond to numbered fractions in the chromatogram above. The amount of material in each lane is normalized to constant absorbance at 280 nm,. The proteins were stained using Flamingo (Biorad).

A

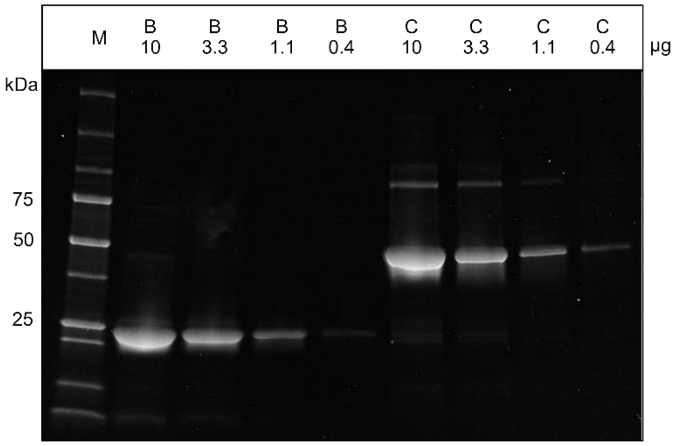


C

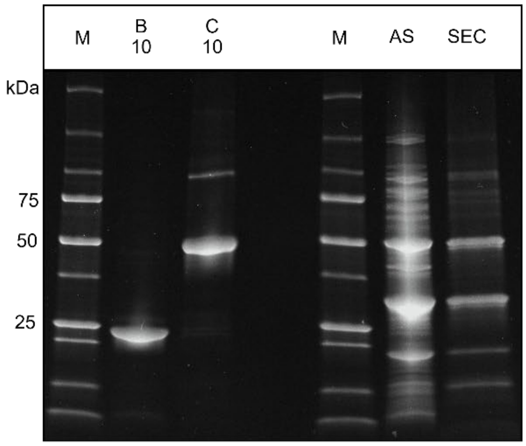


B

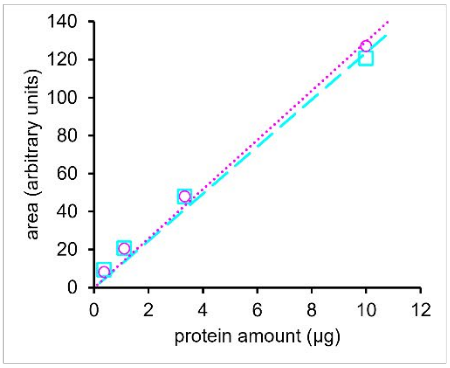


D

| | AccB (lane B) | AccC (lane C) | response ratio B/C |
|---|---|---|---|
| band area | 464 | 484 | 0.96 |

| | AS area | SEC area |
|---|---|---|
| C | 364 | 247 |
| A+D | 655 | 265 |
| B | 240 | 81 |

| ratio | µg ratio AS | µg ratio SEC | molar ratio AS | molar ratio SEC |
|---|---|---|---|---|
| (A+D)/C | 1.80 | 1.08 | 1.30 | 0.77 |
| B/C | 0.66 | 0.33 | 1.95 | 0.97 |

| | AccB/AccC | AccA+D/AccC |
|---|---|---|
| kDa ratio | 0.34 | 1.39 |

Figure S3. Densitometry analysis of the purified ACC complex. (**A**) SDS-PAGE of purified his-tagged AccB and his-tagged AccC stained with flamingo dye. The numbers above the lanes indicate the amount of each protein loaded based on UV/Vis quantification. The 260/280 ratio is 0.70 for AccB and 0.93 for AccC. These gel images have not been contrast or brightness modified, and oversaturation was not detected. (**B**) Quantification of the dye response as a function of protein loading for SDS-PAGE from panel (A). All bands in each lane are used for quantification, with the cyan squares representing AccB and magenta circles representing AccC. The response of AccB is roughly equal to AccC under the conditions used. This analysis confirms a linear response under the conditions used, see methods for recommended procedure used here. Variation in gel fixing, staining and destaining times can alter the response ratio between AccB/AccC. (**C**) SDS-PAGE of the pRSF derived 30% ammonium sulfate pellet in lane AS or SEC fraction, and purified his-tagged AccB in lane B or AccC in lane C. Each lane has 45 µg total protein is loaded. (**D**) Densitometry analysis of panel (C). The lanes B and C are internal controls to determine the response ratio and used to correct for the integration area of AccB. The bands for AccA and AccD (and BirA if present) are integrated together since the peaks significantly overlap. Dividing the corrected area of AccB by the area of AccC gives the ratio in terms of µg. The difference in molecular weight is used to provide the final molar ratio. The molar ratio difference for AccA/D in the SEC fraction reveals there is likely a 20% difference in staining efficiency compared to AccC. Comparison of the AS and SEC bands also indicates a loss of excess AccA/AccD or BirA that co-precipitates with the complex.

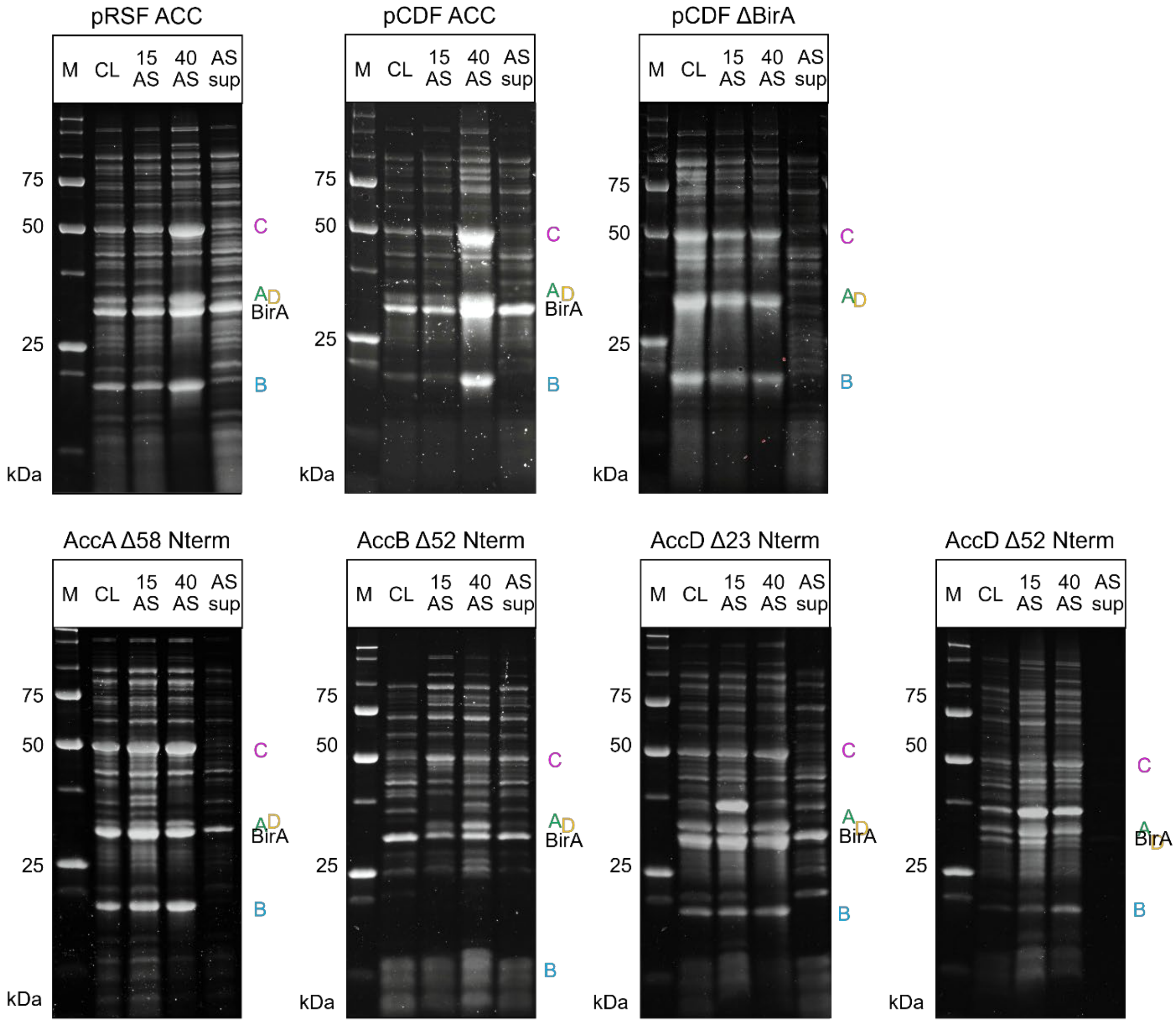


Figure S4. Confirmation of ACC subunit productionfor ACC and N-terminal deletion mutants expressed from pCDF. SDS-PAGE of clarified lysate (CL) and ammonium sulfate (AS) fractions, either 15% pellet, 40% pellet or 40% supernatant (sup). The gels were stained using Flamingo (Biorad).

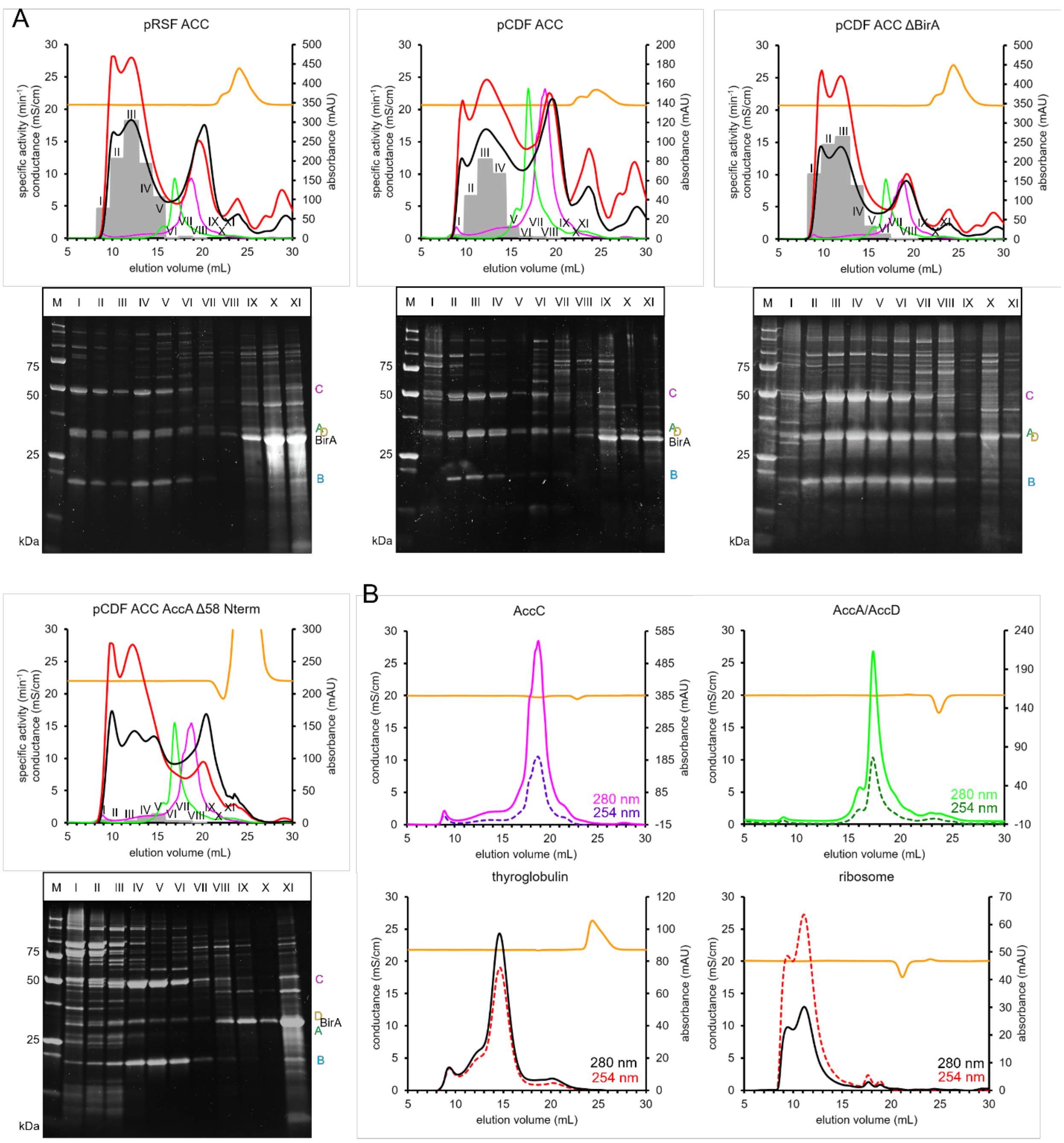


Figure S5. SEC purification of the ACC complexes from pCDF. A) Top panels - chromatograms are from a Superose 6 Increase 10/300 GL column. Absorbance at 280 nm as black lines and 260 nm as red lines. The conductance is shown as an orange line. The gray bars indicate the specific activity for a set of pooled fractions (roman numerals) normalized to a constant protein concentration determined by a ratiometric bradford assay. Bottom panels - SDS-PAGE for the fractions after Flamingo staining. Similar total amounts of protein are loaded. Normalized traces for AccC and AccA/D at 280 nm are shown as magenta and green lines respectively. B) Traces for AccA/D (~135 kDa), AccC (~99 kDa), thyroglobulin (~660 kDa) and the *E. coli* 70S ribosome for comparison to the ACC complex purification.

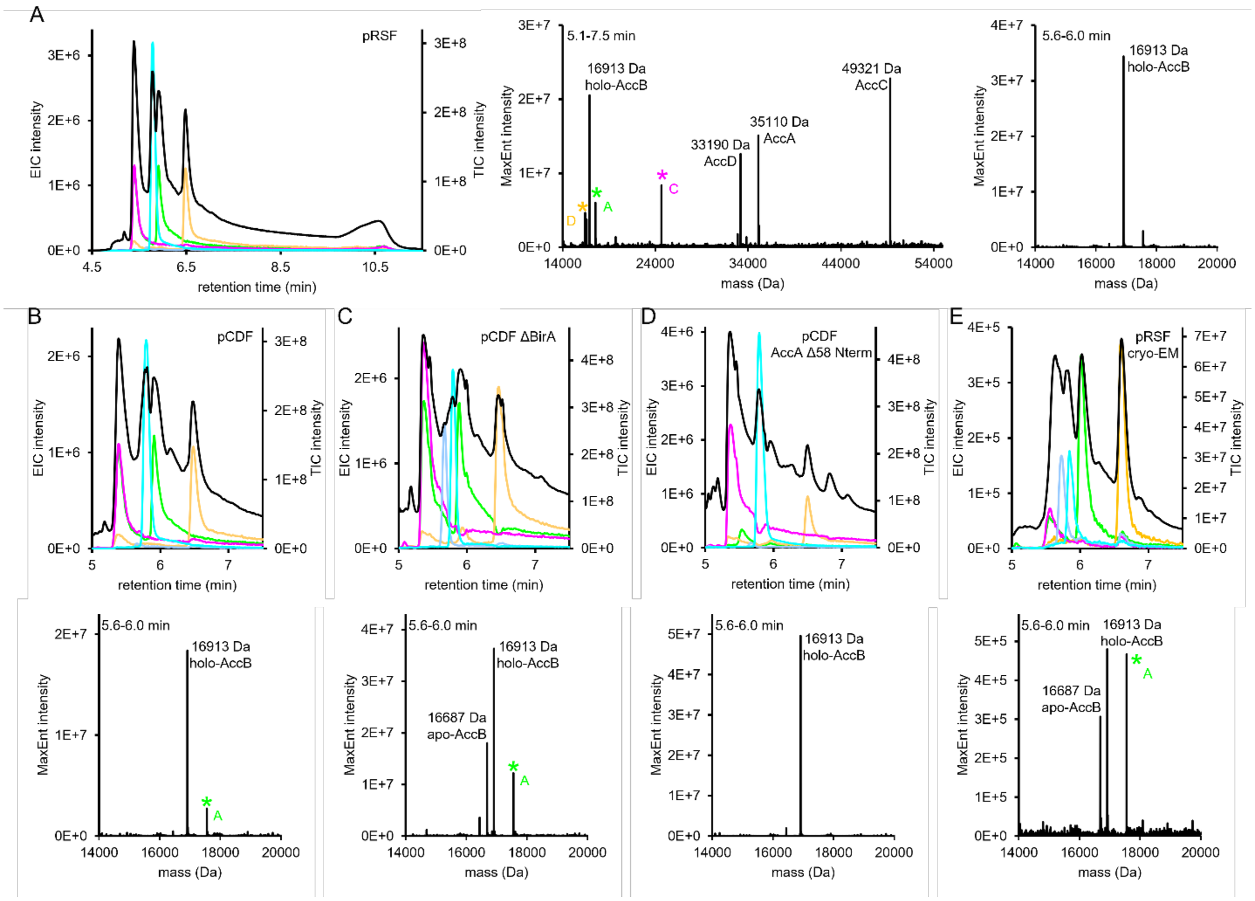


Figure S6. Intact protein mass spectrometry. (**A**) Left panel – chromatogram of ACC from the pRSF plasmid, black line is the total ion chromatogram (TIC) and extracted ion chromatogram (EIC) for AccA represented by m/z 836.9±0.1 in green, AccD 852.0±0.1 in orange, AccC 809.6±0.1 in magenta, apo-AccB 982.5±0.1 in periwinkle and holo-AccB 995.8±0.1 in cyan (this color scheme is consistent for the chromatograms of figures B-E). Middle panel – MaxEnt spectra for a time window that includes all the subunits, 5.1-5.7 min. Note that ½ mass peaks are apparent and are designated by asterisks. Right panel – MaxEnt spectra for a time window that only includes apo- and holo-AccB, 5.6-6.0 min. EIC/TIC and MaxEnt spectra for (**B**) ACC from pCDF, (**C**) ACC from pCDF ΔBirA, (**D**) ACC from pCDF AccA Δ58 N-term and (**E**) ACC from pRSF that was used for cryo-EM.

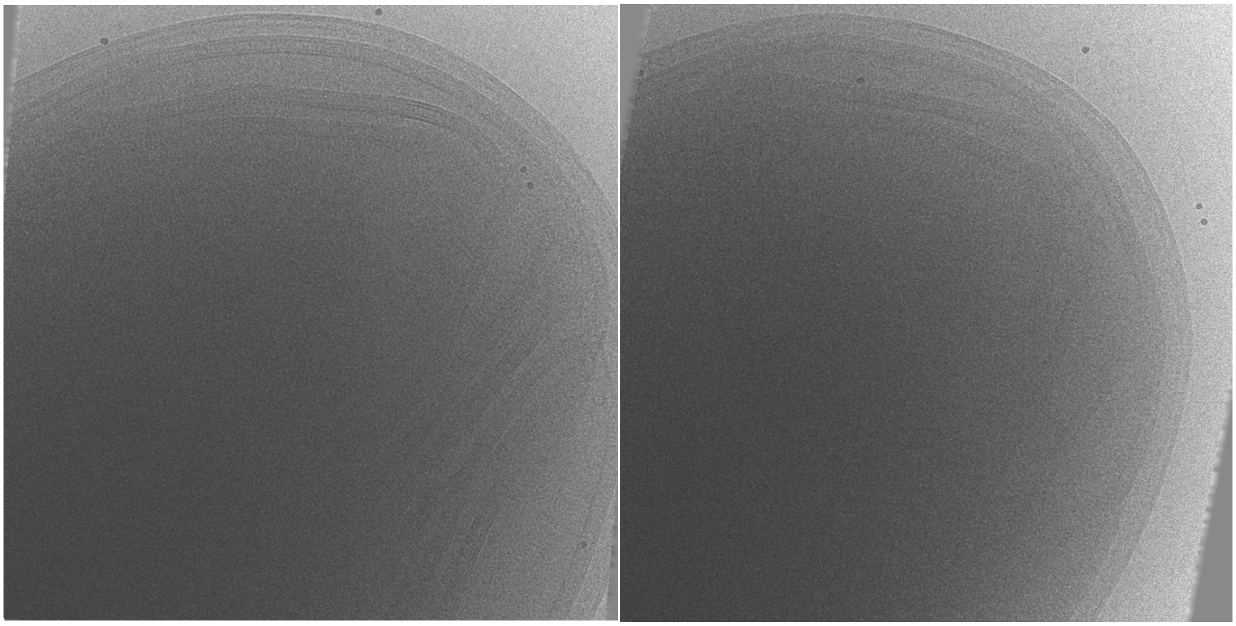

Figure S7. Cryo-EM of intact *E. coli* BL21(DE3) expressing the ACC complex from pRSF. Note the long tubes. Movies of the tilt series are included in the supplementary materials.

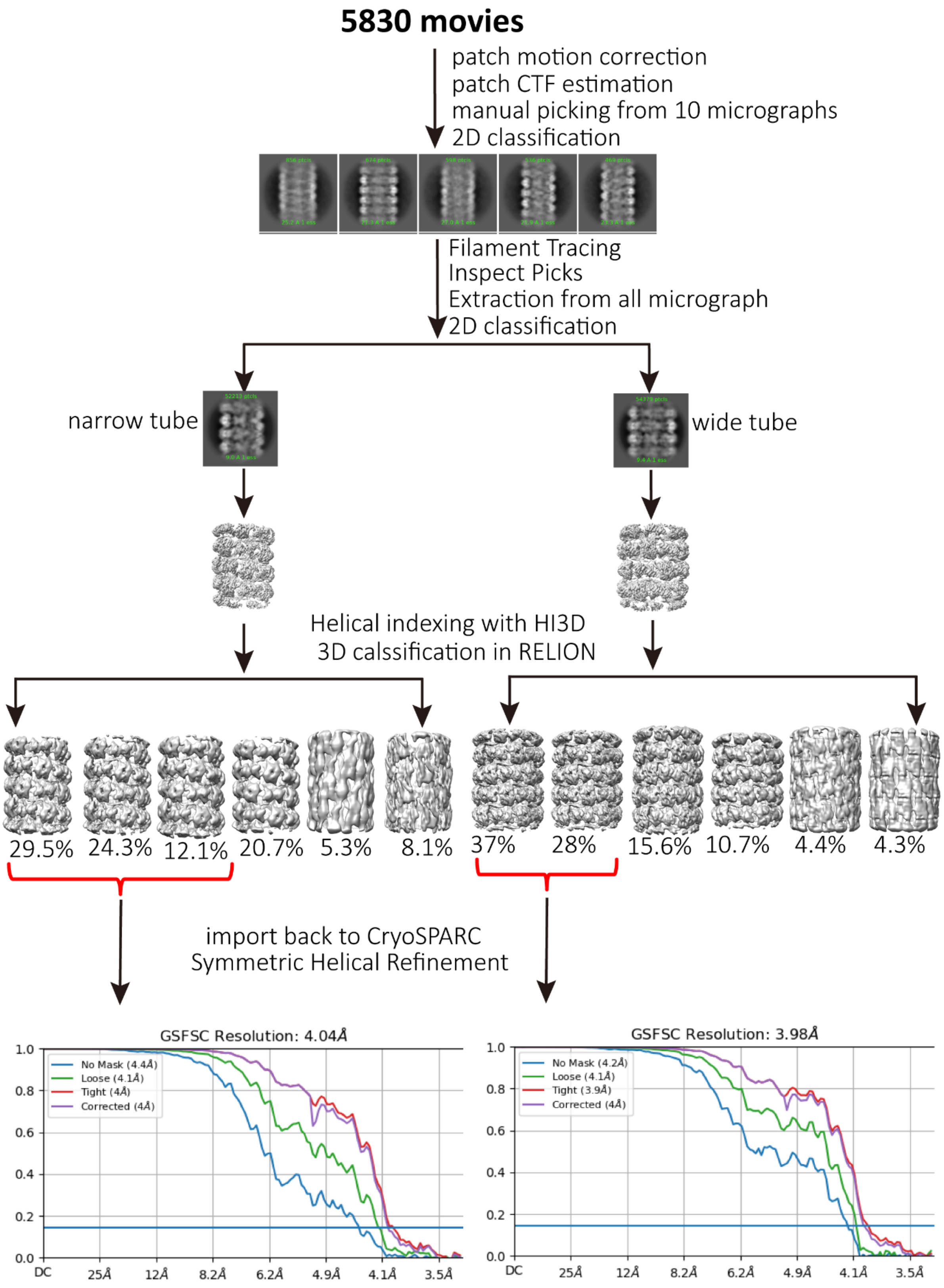


Figure S8. Image processing workflow for narrow and wide tubes of *E. coli* ACC. FSC curves for the two tubes are shown at the bottom.

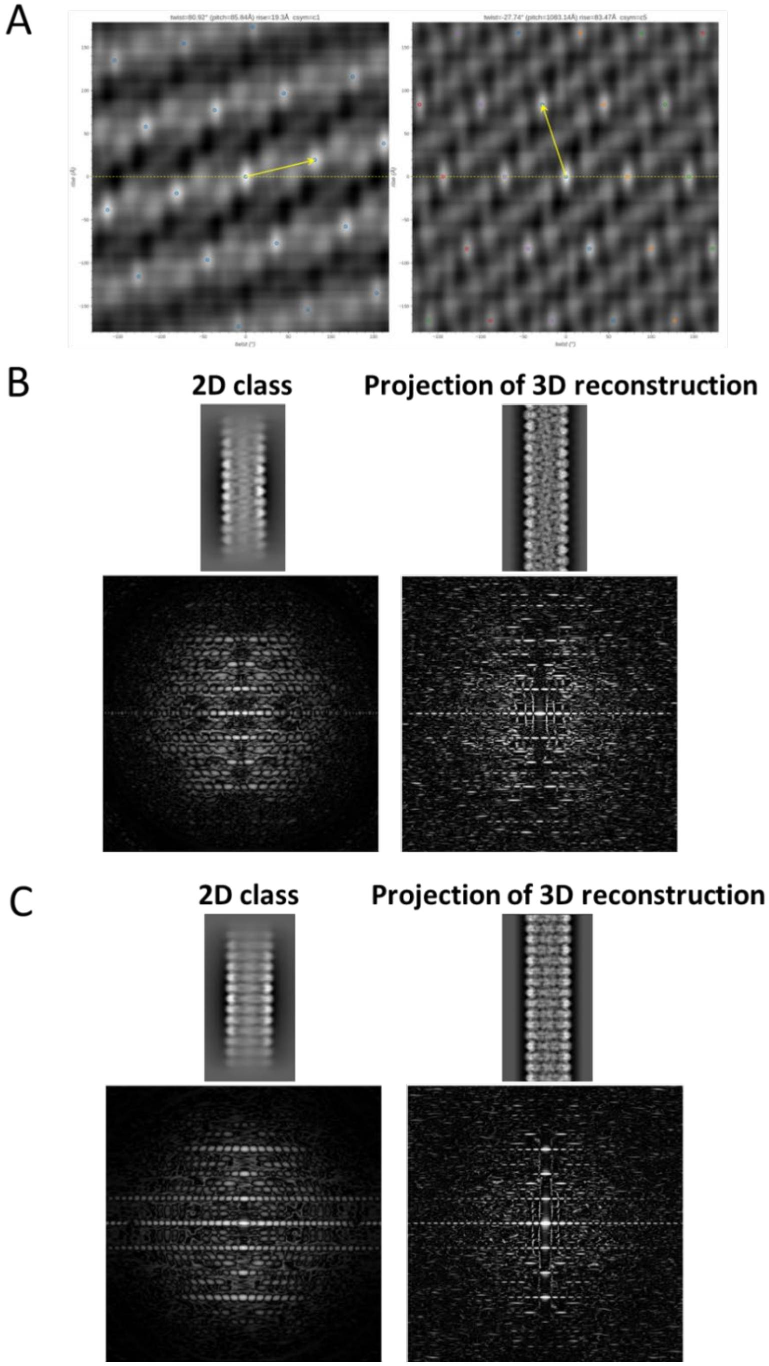


Figure S9. Helical indexing of *E. coli* ACC tubes. (**A**) helical indexing of narrow tube (left) and wide tube (right) in real space using HI3D. The unit cell vectors (yellow arrows) with the shortest distances to the equators correspond to the helical twists (x-coordinate) and rises (y-coordinate) of narrow and wide tubes, respectively. (**B** and **C**) 2D class averages (top, left), back projections from the 3D reconstructions (top, right) and their corresponding power spectra (below) of narrow tube and wide tube. The power spectra were generated using the HILL Web App developed by the Wen Jiang lab (https://jianglab.science.psu.edu/hill). The power spectra were truncated at 15 Å.

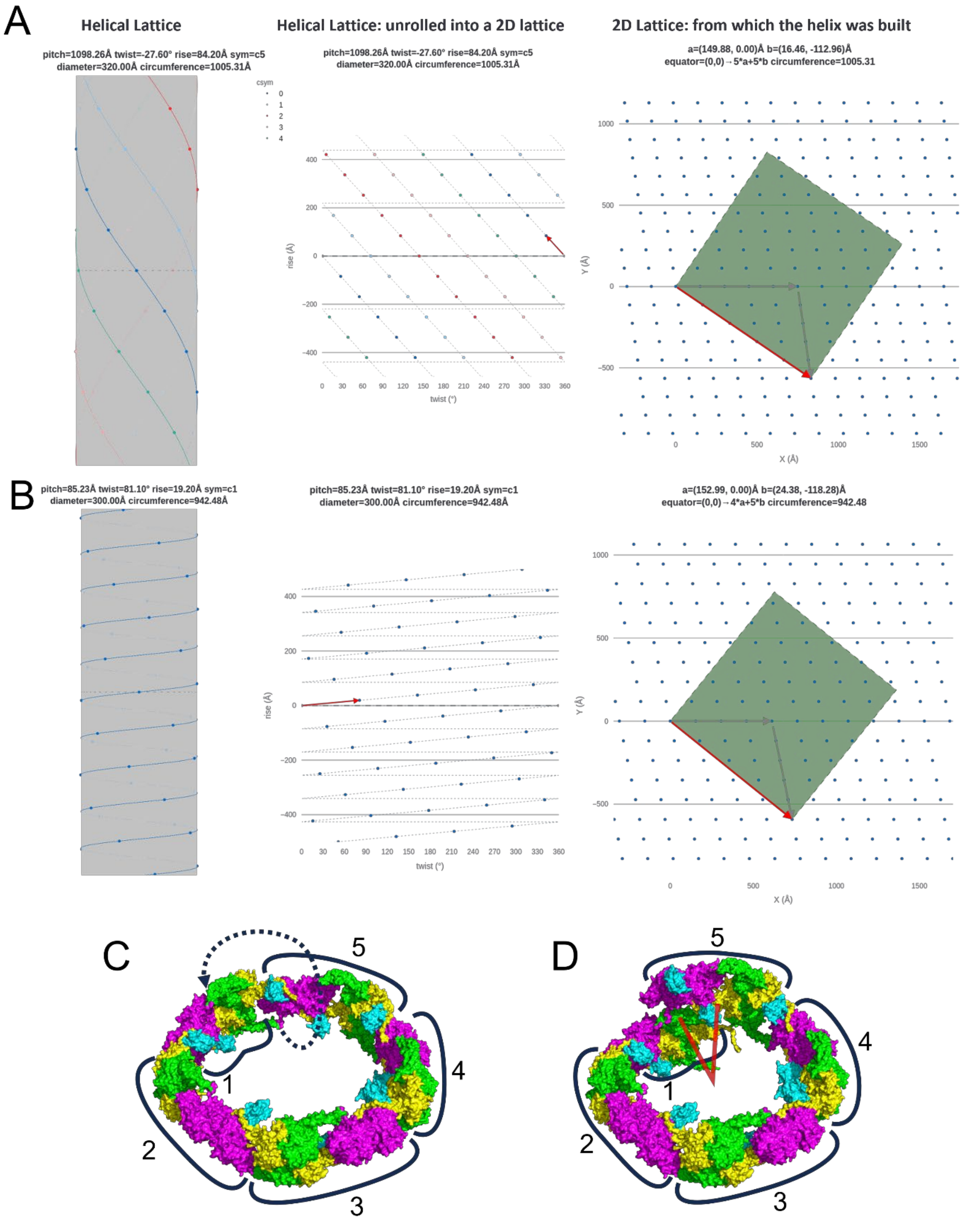


Figure S10. Comparison of tube lattices and single rings. Lattice schematics from https://jianglab.science.psu.edu/helicallattice for **(A)** wide tubes and (**B**) narrow tubes. (**C**) wide tube resembles a single ring. (**D**) narrow tube is helical in nature. Both systems are shown with an equivalent number of subunits. Moving protomer 5 up and over from the annular system generates the helical system, as shown with a dashed line.

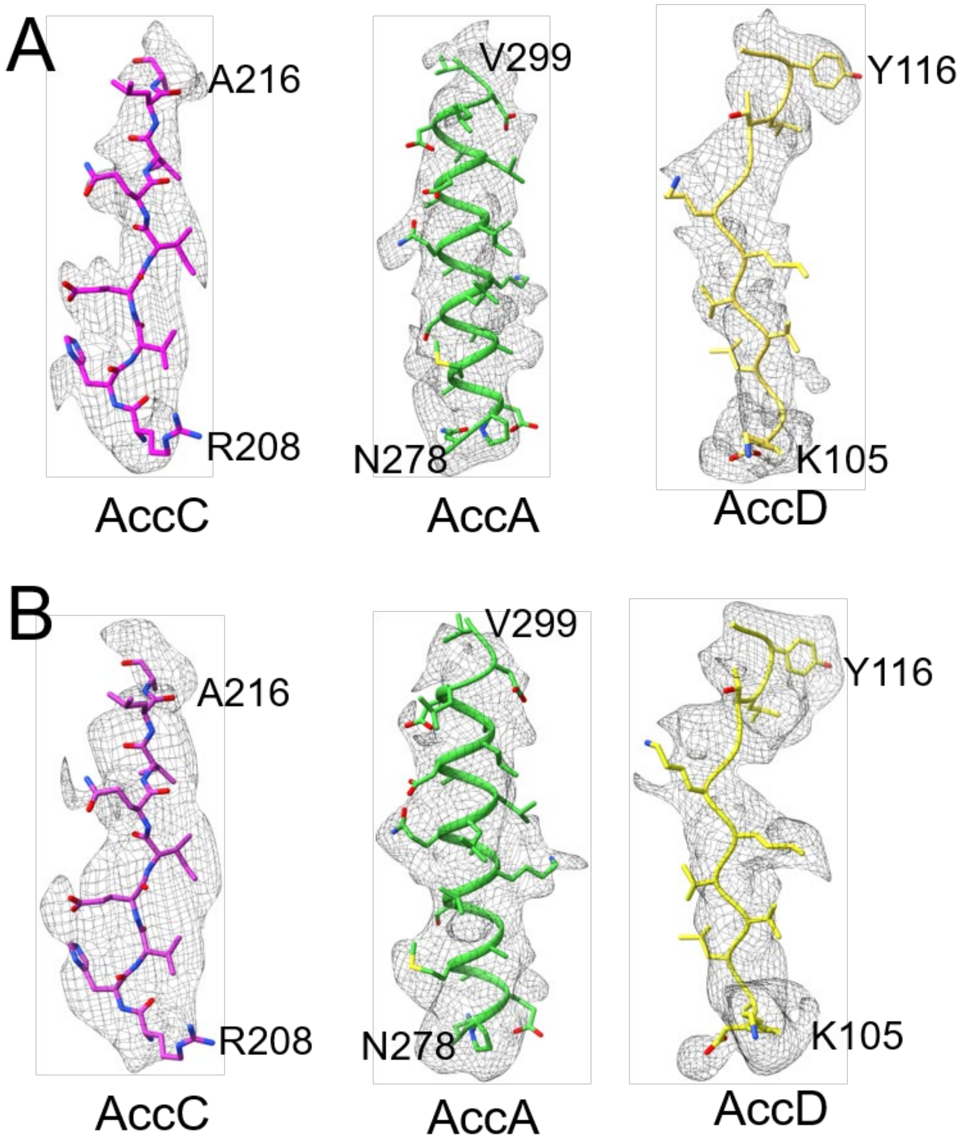


Figure S11. Cryo-EM map examples of initial helical reconstructions at ~4.0 Å overall resolutions. (**A** and **B**) Representative regions of the cryo-EM maps superimposed on the atomic models of the narrow and wide tubes, respectively. The corresponding regions are also shown to demonstrate the improved quality of subsequent symmetry-expanded and locally refined maps shown in Fig. S13E and S13J.

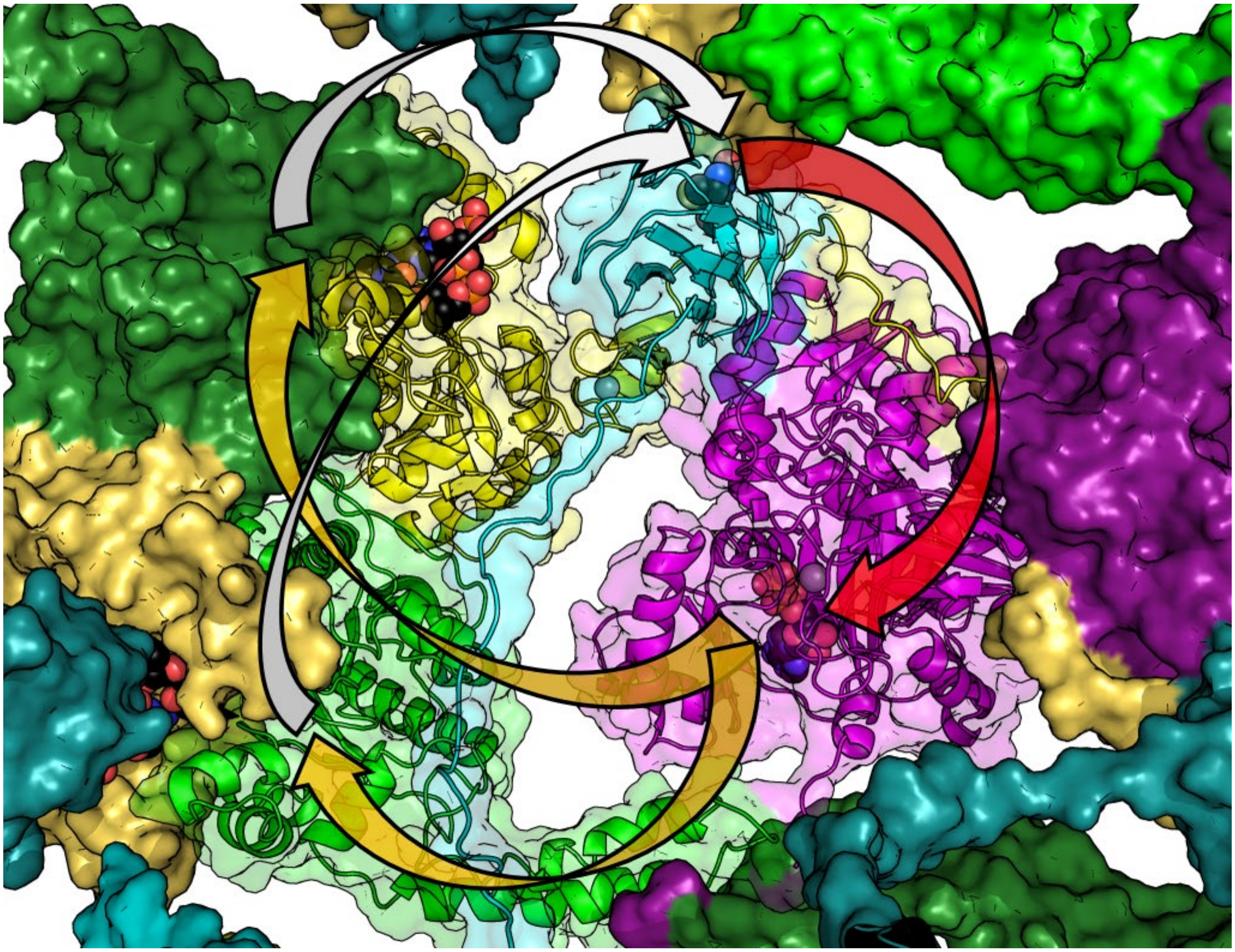

Figure S12. Overall structure of one heteromeric unit and orientation of the active sites. Transparent surface demonstrates the close positioning of the BCCP docking site and catalytic active sites. The arrows represent movement of the BCCP from the docking site to the BC active site (red arrow), to the CT active sites (orange arrows) and back to the docking site (white arrows), representing one catalytic cycle.

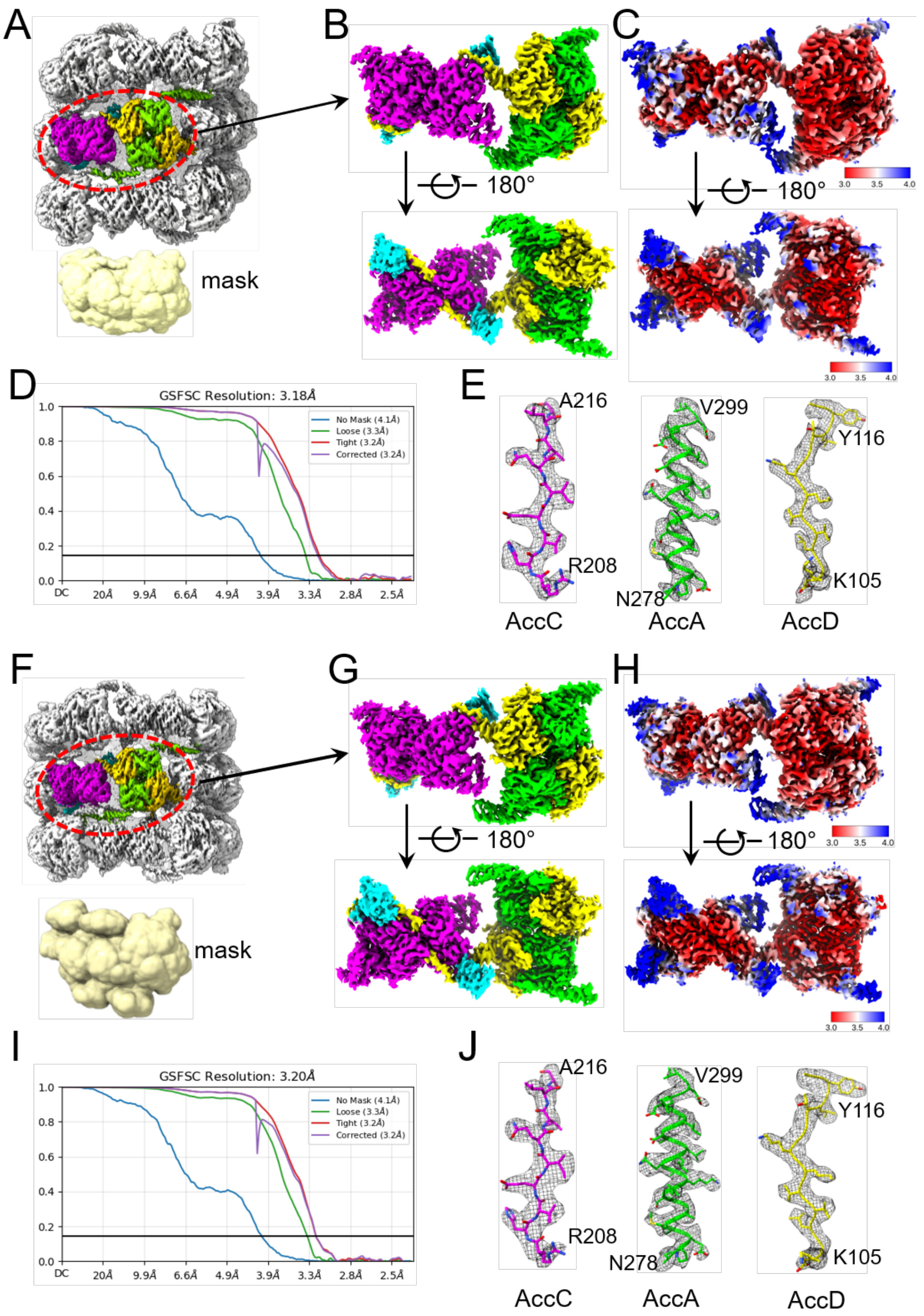


Figure S13. Local reconstructions of *E. coli* ACC tubes. (**A** and **F**) The masked regions (upper) and the masks (lower) used for local refinement of the narrow and wide tubes, respectively. (**B** and **G**) The density maps of the narrow and wide tubes after local reconstruction, respectively. The color scheme of BC, BCCP and CT is the same as shown in Fig. 1. (**C** and **H**) The same maps as shown in (B and G), but the colors represent the local resolutions. (**D** and **I**) FSC curves for the local reconstructions of the narrow and wide tubes, respectively. (**E** and **J**) Parts of the cryo-EM electron potential maps superimposed upon the atomic models of narrow and wide tubes, respectively.

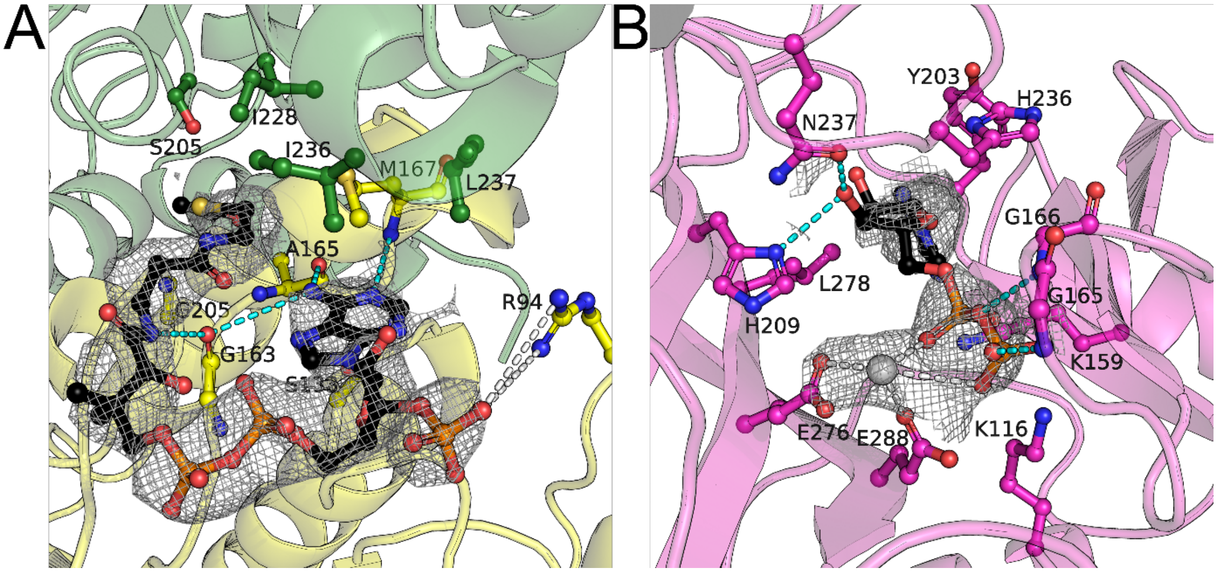


Figure S14. Ligand density and interactions. (**A**) CoA/acetyl-CoA density and interactions with the CT domain. (**B**) ADP density and interactions with the BC domain. Putative hydrogen bonds are shown as cyan dashes and electrostatic interactions are white dashes.

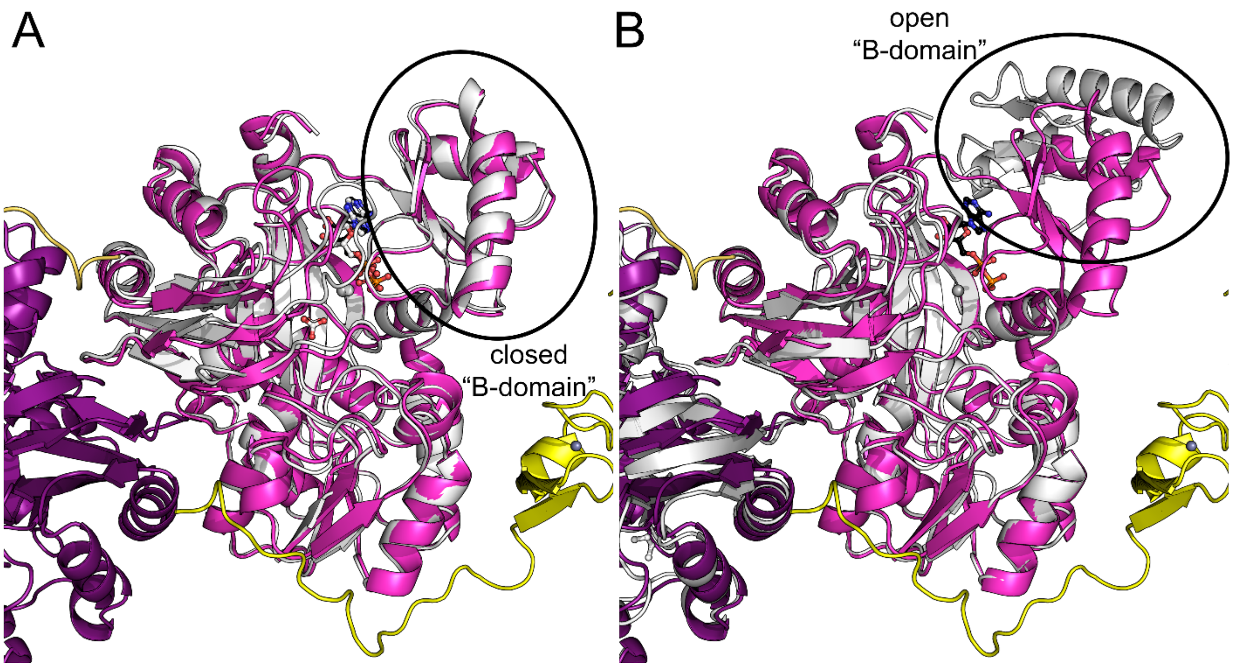


Figure S15. BC domain conformation comparisons. Comparison of our cryo-EM structure showing the BC domain in magenta and purple for the dimeric mate with (**A**) PDB 3RV4 in light gray or (**B**) PDB 4HR7 chain A in light gray. The N-terminus and zinc-finger domain of AccD is shown in yellow for reference.

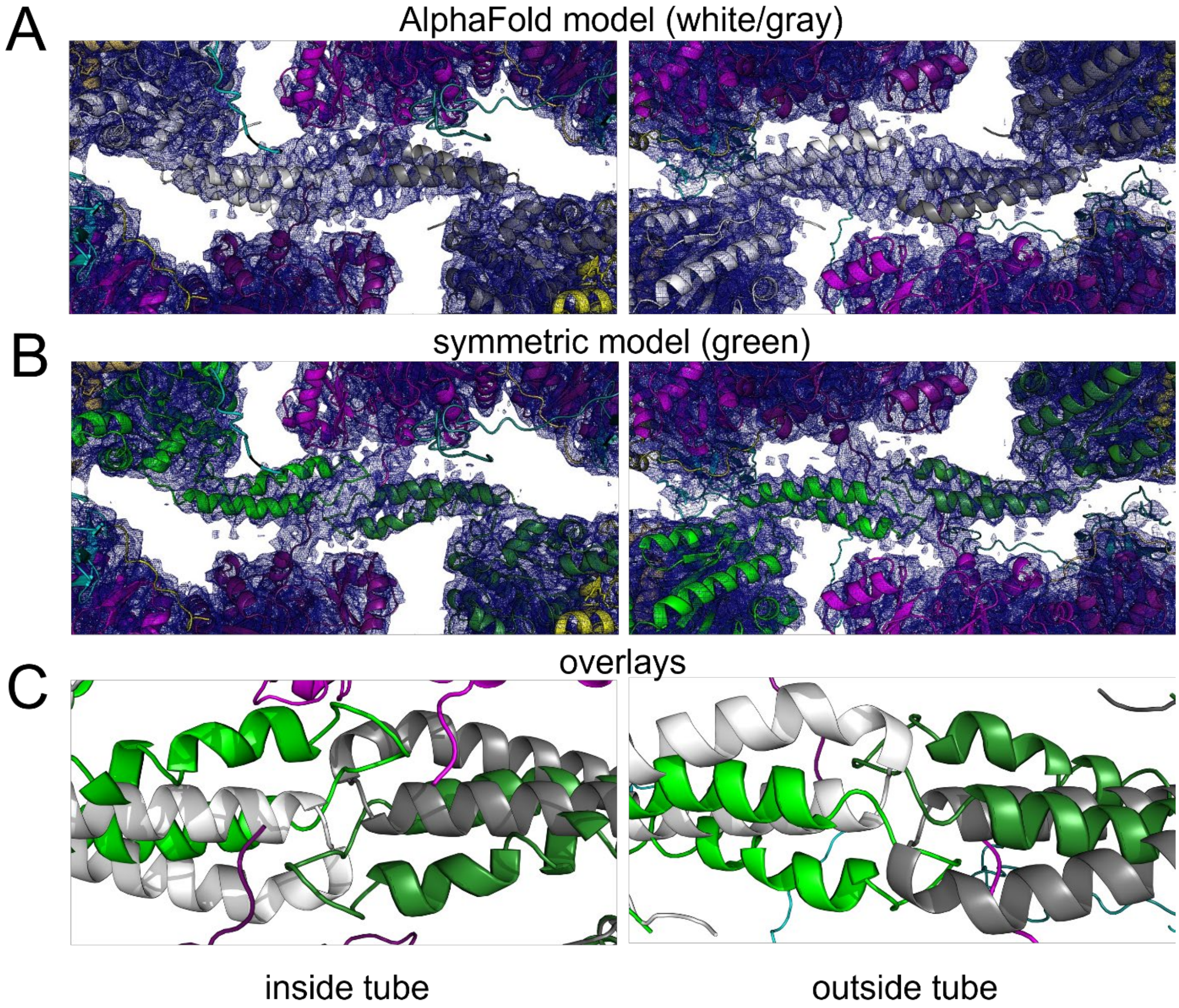


Figure S16. AccA N-terminal helix interaction cryo-EM density. (**A**) The main body of the AccA AlphaFold model was fit into the map, AccA shown in white and gray. (**B**) One model for how the AccA N-terminus might interact, AccA shown in green and dark green. Notice that the C-terminal tails of the BC (magenta) are fit into density that is continuous with the AccA N-terminal helices. (**C**) Comparison of the AlphaFold model and refined symmetrical model.

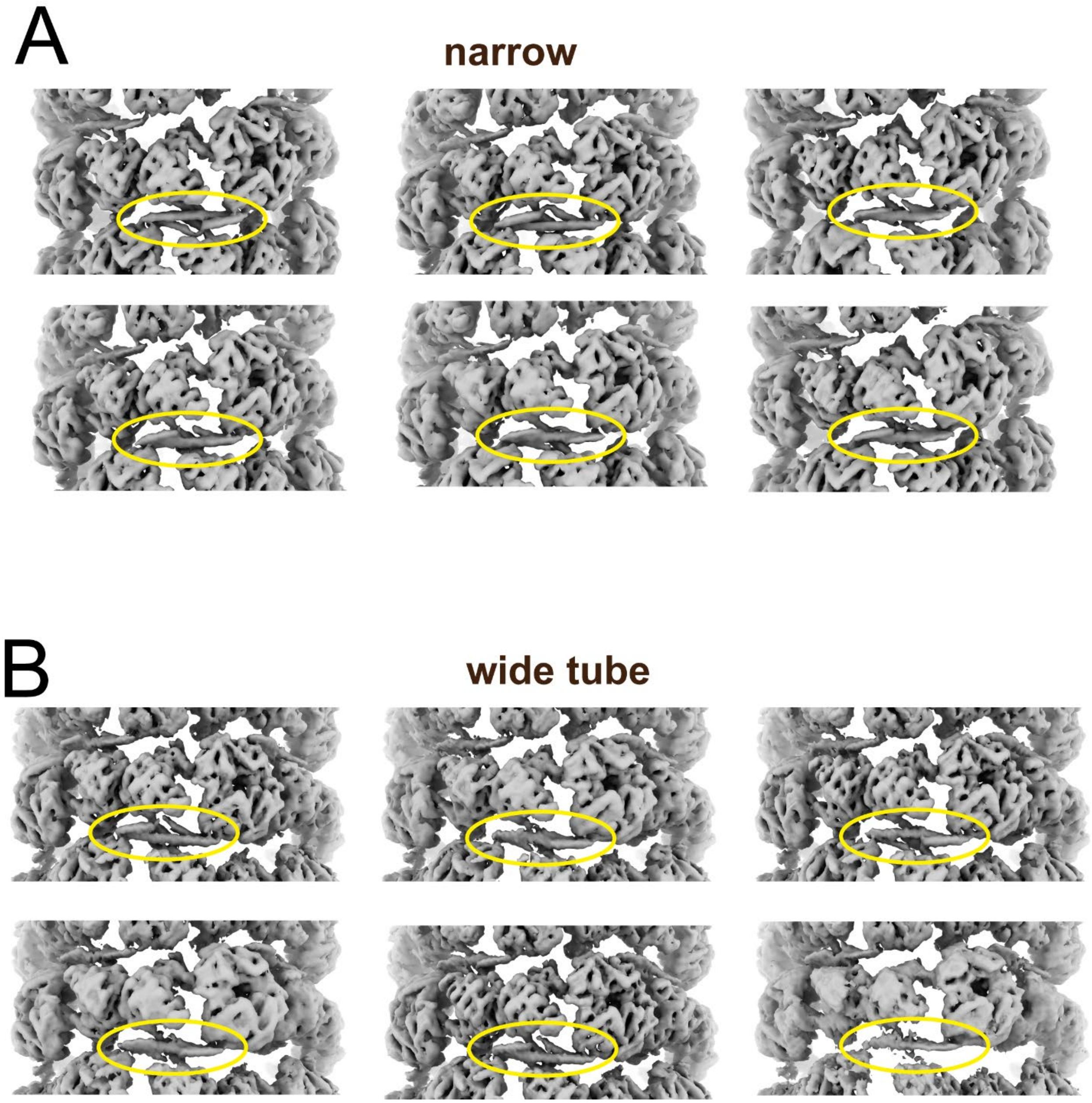


Figure S17. Local 3D classification and reconstructions of regions around the AccA N-terminal helices for the (**A**) narrow tubes and (**B**) wide tubes.

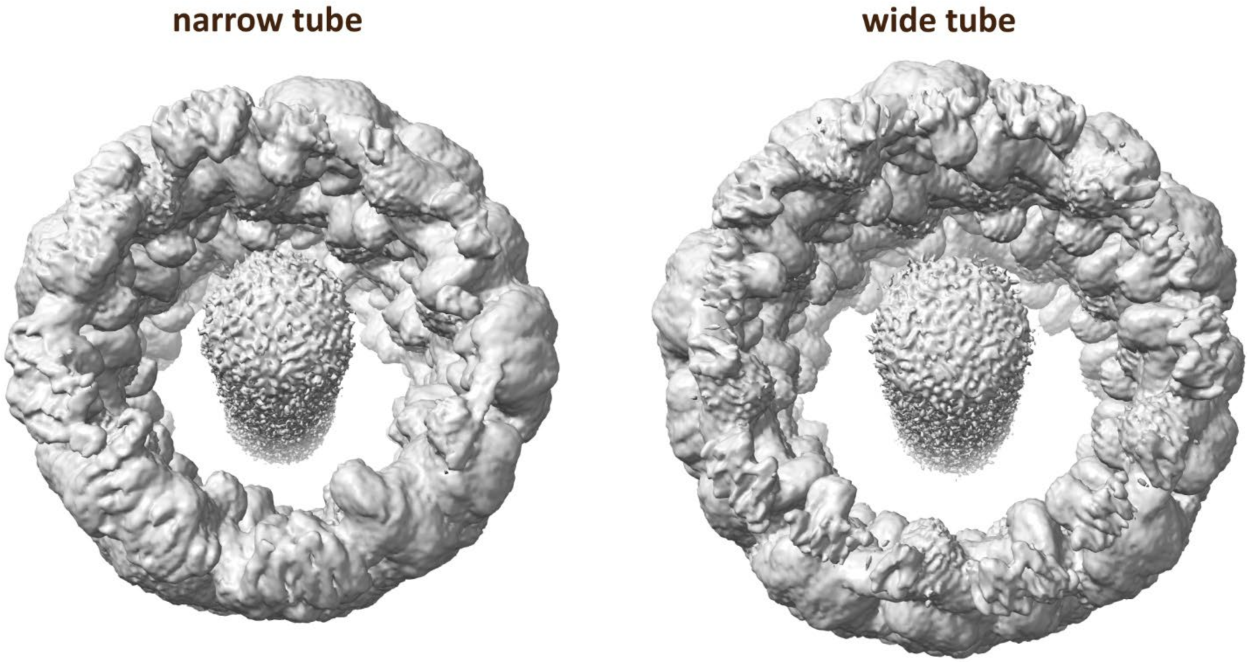


Figure S18. Densities along the helical axis in the two tube structures.

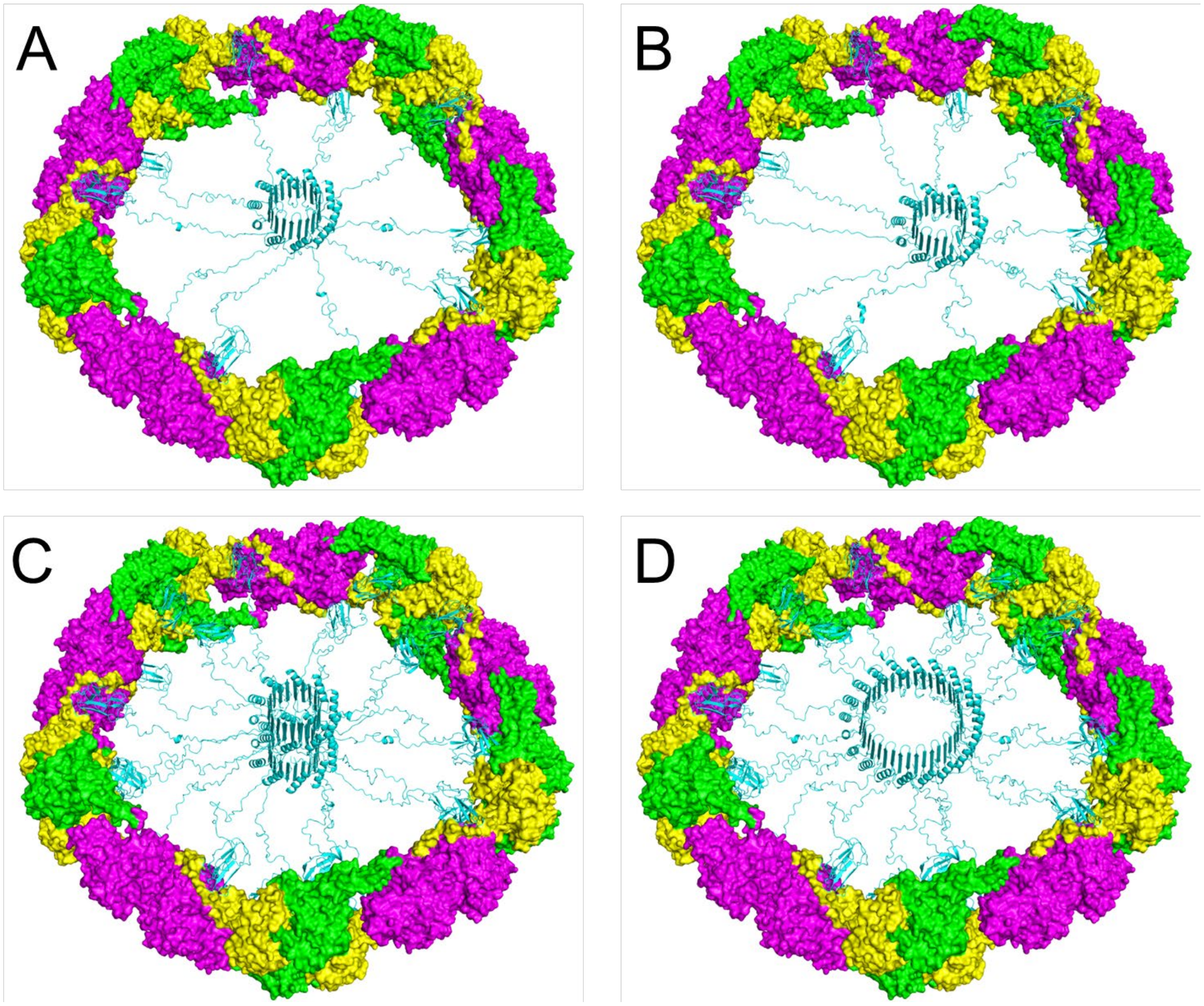


Figure S19. Alternative models for interactions of the AccB N-termini. (**A**) Model of 10 AccB N-termini that are centrally located in a single wide tube ring. (**B**) Similar to (A) but offset. The offset location reveals there is significant flexibility in the location of the putative β-barrel. (**C**) Two sets of AccB in a single wide tube ring. The second copy of the AccB C-terminal domain in each protomer is placed at the putative entrance to the acetyl-CoA binding site. (**D**) Similar to (C) but the 20 AccB N-termini are modeled as a large beta barrel. While the current biochemical data suggests protomers have a 1:1:1 BC:BCCP:CT ratio, there is space available in the ring to accommodate twice the number of BCCPs. These models are available at doi.org/10.5281/zenodo.14743899

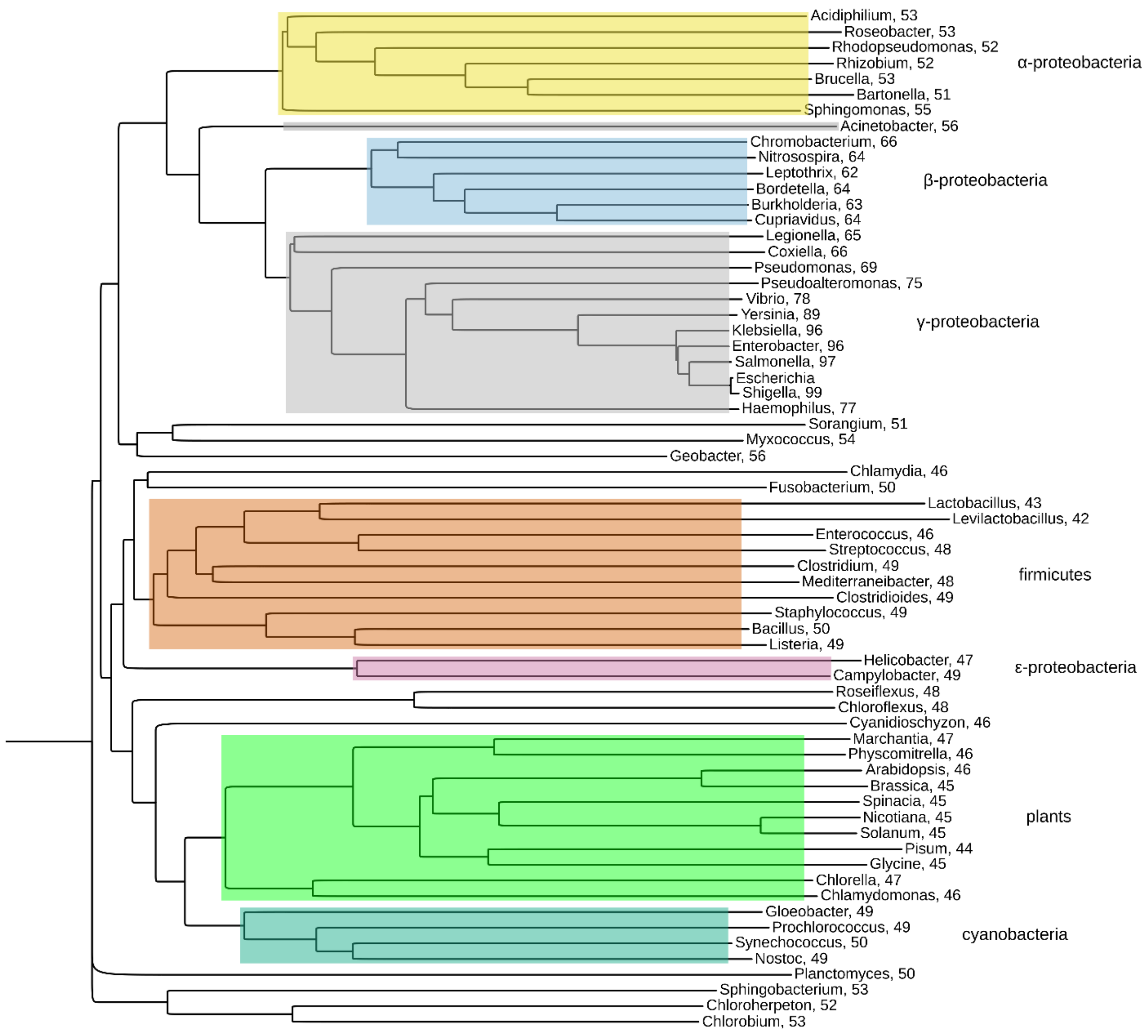


Figure S20. Phylogenetic tree of concatenated AccC/B/D/A homologs. Sequences were aligned and tree generated with Clustal Omega. The overall sequence identity percentage to *E. coli* is shown after the genus name. The accession codes used to generate the sequence alignment and tree are as follows:

Escherichia NP_417722.1/NP_417721.1/NP_416819.1/NP_414727.1
Shigella NP_709051.2/NP_709050.1/NP_708198.2/NP_706130.1
Salmonella WP_000884627.1/WP_000354626.1/WP_000118383.1/WP_000055753.1
Enterobacter WP_006178859.1/WP_008502856.1/WP_008502598.1/WP_008501895.1
Klebsiella WP_004174096.1/WP_004206250.1/WP_004201773.1/WP_004145855.1
Yersinia WP_002210069.1/WP_002230506.1/WP_002209729.1/WP_002212147.1
Vibrio WP_000884888.1/WP_000354612.1/WP_000118592.1/WP_002032753.1
Haemophilus WP_005669669.1/WP_006995619.1/WP_006995883.1/WP_005633993.1
Pseudoalteromonas WP_010562121.1/WP_010562120.1/WP_010559658.1/WP_010559708.1
Pseudomonas WP_003095391.1/WP_003110495.1/WP_003104199.1/WP_003109333.1
Coxiella NP_820707.1/NP_820706.1/NP_819908.1/NP_820493.1
Legionella WP_011213101.1/WP_010946212.1/WP_010947071.1/WP_010946522.1
Nitrosospira WP_011382031.1/WP_011382032.1/WP_011381225.1/WP_011381127.1
Chromobacterium WP_011134540.1/WP_081526683.1/WP_011136307.1/WP_011136737.1

Cupriavidus WP_012353874.1/WP_012353873.1/WP_012353352.1/WP_012352493.1
Burkholderia WP_004194420.1/WP_004194067.1/WP_004188147.1/WP_004193249.1
Bordetella WP_003814951.1/WP_003814952.1/WP_003813843.1/WP_003810007.1
Leptothrix WP_012348985.1/WP_012348984.1/WP_012346712.1/WP_012346700.1
Acinetobacter WP_005285498.1/WP_005285506.1/WP_005309656.1/WP_005228791.1
Sphingomonas WP_007405803.1/WP_007405762.1/WP_007403603.1/WP_007403457.1
Bartonella WP_011180701.1/WP_011180702.1/WP_038486627.1/WP_011181396.1
Brucella WP_002966788.1/WP_002964036.1/WP_002965171.1/WP_002965096.1
Rhizobium WP_018241835.1/WP_047622256.1/WP_018068910.1/WP_018448126.1
Rhodopseudomonas WP_011157985.1/WP_011157986.1/WP_011155642.1/WP_011156076.1
Roseobacter WP_011568993.1/WP_011568992.1/WP_011566674.1/WP_011567285.1
Acidiphilium WP_172637330.1/WP_013640790.1/WP_007422854.1/WP_007423469.1
Sorangium WP_012236552.1/WP_157906787.1/WP_012238414.1/WP_012240223.1
Myxococcus WP_011555718.1/WP_011555719.1/WP_011552772.1/WP_011554047.1
Geobacter WP_010942662.1/WP_010942663.1/WP_010943005.1/WP_010942051.1
Sphingobacterium WP_149913112.1/WP_120333456.1/WP_159730536.1/WP_120259503.1
Chloroherpeton WP_012499260.1/WP_012499259.1/WP_012499972.1/WP_157452565.1
Chlorobium WP_011890836.1/WP_126341565.1/WP_126341739.1/WP_011890832.1
Planctomyces WP_013108347.1/WP_013108348.1/WP_013108928.1/WP_013110431.1
Chloroflexus WP_012259259.1/WP_242604991.1/WP_012257520.1/WP_012257519.1
Roseiflexus WP_012121417.1/WP_012121385.1/WP_041330882.1/WP_012121414.1
Nostoc WP_012412185.1/WP_012407244.1/WP_012409325.1/WP_012408398.1
Synechococcus WP_011242491.1/WP_011243856.1/WP_011244449.1/WP_011242365.1
Prochlorococcus WP_011130955.1/WP_011129411.1/WP_011129913.1/WP_011130598.1
Gloeobacter WP_011142010.1/WP_011141286.1/WP_011141604.1/WP_011142858.1
Cyanidioschyzon XP_005538926.1/NP_849043.1/NP_849109.1/NP_848979.1
Campylobacter YP_002344680.1/YP_002344681.1/YP_002343587.1/YP_002343880.1
Helicobacter WP_001031226.1/WP_001053821.1/WP_000505081.1/WP_001029379.1
Fusobacterium WP_005979325.1/WP_005982233.1/WP_005981108.1/WP_005981110.1
Chlamydia WP_010882833.1/WP_010882834.1/WP_010882708.1/WP_010883057.1
Listeria NP_464882.1/NP_464881.1/NP_465098.1/NP_465097.1
Bacillus WP_003230253.1/WP_003230252.1/WP_003223544.1/WP_003229417.1
Staphylococcus WP_000756612.1/WP_001009516.1/WP_000471568.1/WP_000883645.1
Clostridioides WP_003435334.1/WP_003428740.1/WP_003424064.1/WP_009889903.1
Mediterraneibacter WP_020436402.1/WP_015528346.1/ /WP_055146213.1
Clostridium WP_011098458.1/WP_011098456.1/ /WP_011098459.1
Streptococcus WP_000488675.1/WP_001052244.1/WP_001173353.1/WP_001017399.1
Enterococcus WP_002295381.1/WP_002356285.1/WP_002290846.1/WP_002362363.1
Levilactobacillus WP_042516774.1/WP_021742581.1/WP_039104729.1/WP_015473665.1
Lactobacillus WP_127345880.1/WP_080881990.1/WP_127345879.1/WP_080881986.1
Arabidopsis NP_198386.1/NP_197143.1/NP_051068.1/NP_565880.1
Brassica XP_013716504.1/XP_022567554.1/YP_005089961.1/XP_048631667.1
Glycine NP_001237138.1/NP_001238375.1/YP_538774.1/NP_001235234.2
Nicotiana NP_001312641.1/NP_001312387.1/NP_054508.1/XP_016462645.1
Pisum XP_050893622.1/XP_050911986.1/AXY95782.1/Q41008.1
Solanum XP_006348560.1/XP_006345777.1/YP_009915983.1/XP_006363070.1
Spinacia XP_021867376.1/XP_021839458.1/NP_054945.1/XP_021859278.2
Marchantia OAE29968.1/PTQ33718.1/P12217.1/PTQ34056.1
Physcomitrella XP_024381777.1/XP_024397898.1/Q00761.1/XP_024356681.1
Chlorella KAI3436743.1/KAI3428383.1/NP_045833.1/KAI3431329.1
Chlamydomonas XP_042921783.1/XP_001700442.1/XP_001703187.2/XP_001696945.1

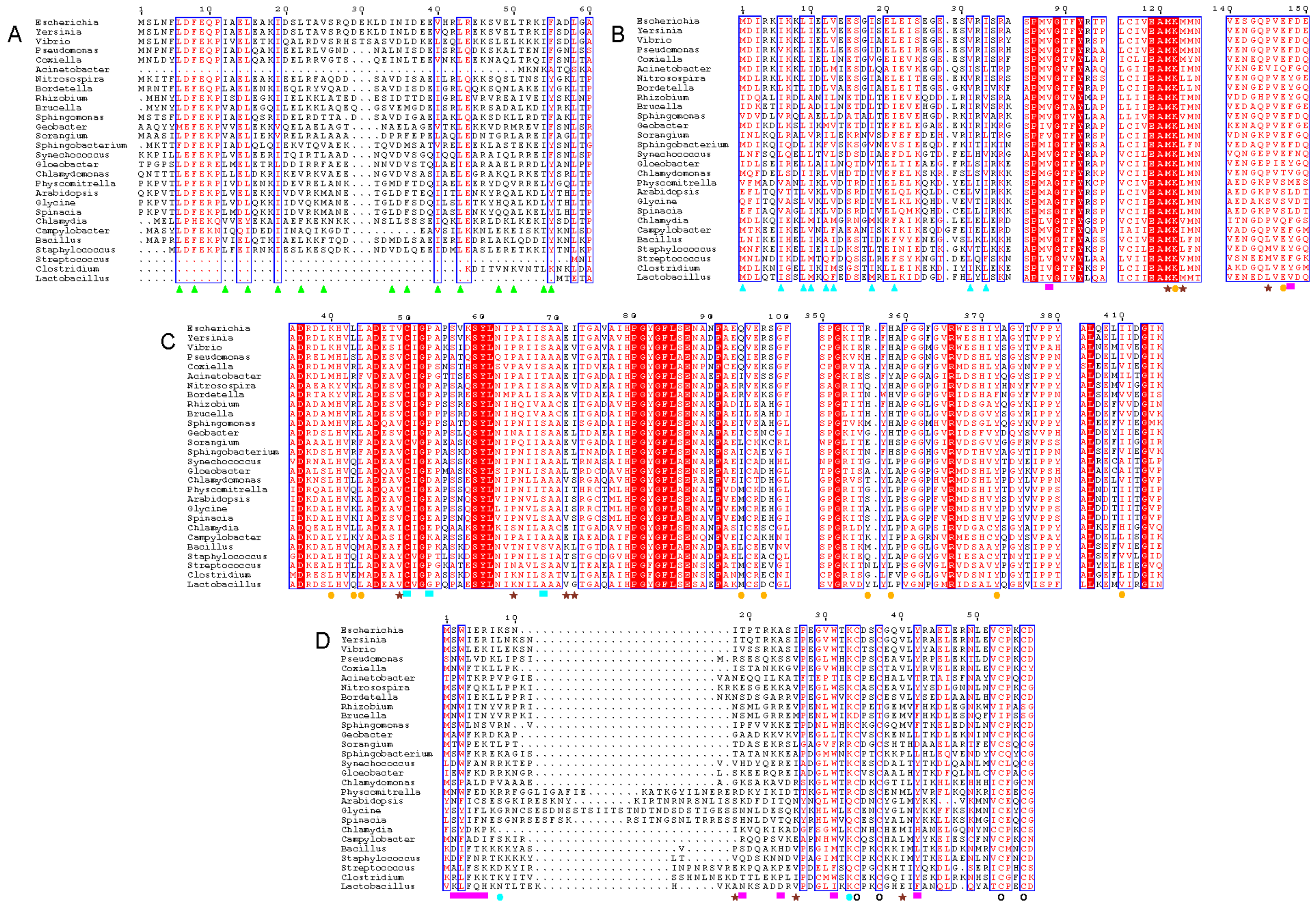

Figure S21. Protein:protein interaction homology. Portions of sequence alignments annotated with interactions, cyan squares represents AccB interactions, orange circles represents AccD interactions, maroon stars represents two interactions (AccC with B/D, AccB with C/D, and AccD with B/C), green trianges represents hydrophobic interactions between AccA N-terminal helices, cyan triangles represents hydrophobic interactions between N-terminal α/β/β motifs. Zinc-finger cysteines are indicated by open circles.

Supplementary Table 1. ACC complex activity

| sample | Crude lysate specific activity | 30/40% AS pellet specific activity | SEC specific activity | Estimated holo-AccB % |
|---|---|---|---|---|
| pRSF AccA/D/B/C/BirA Superose 6 10/300 GL for cryo-EM | 0.012 | 0.93 | 0.93 | 60 |
| pRSF AccA/D/B/C/BirA Sephacryl S-500 | -- | -- | 5.0 | -- |
| pRSF AccA/D/B/C/BirA* | 0.014 | 0.056 | 0.13 | 100 |
| pCDF AccA/D/B/C/BirA* | 0.023 | 0.035 | 0.09 | 100 |
| pCDF ΔbirA* | 0.022 | 0.040 | 0.12 | 70 |
| pCDF AccA Δ58 N-term* | 0.0064 | 0.010 | 0.01 | 100 |
| pCDF AccB Δ58 N-term* | 0.00037 | ND | -- | -- |
| pCDF AccD Δ23 N-term* | 0.00019 | ND | -- | -- |
| pCDF AccD Δ52 N-term* | 0.00022 | ND | -- | -- |

Specific activity: μmol/min/mg enzyme

ND: not detected

* for direct comparison, growth and purification conditions are identical with protein concentration determined by Bradford assay and where applicable SEC over a Superose 6 Increase 10/300 GL column.

Supplementary Table 2. Cryo-EM data collection and model statistics.

| **Data collection** | |
|---|---|
| Electron microscope | Titan Krios G4 |
| Camera | K3 |
| Magnification | 105000 |
| Voltage (kV) | 300 |
| Defocus range (µm)[a] | 0.8-2.0 |
| Pixel size (Å) | 0.411 (super-resolution pixels), 0.822 (physical pixels) |
| Number of movies | 5,838 |
| Frame exposure time (s) | 2.01 |
| Number of frames | 50 |
| Total electron dose (e/Å$^2$) | 54.438 |

| **Reconstruction** | **Narrow helical tube** | | **Wide helical tube** | |
|---|---|---|---|---|
| | **Helical reconstruction** | **Local reconstruction** | **Helical reconstruction** | **Local reconstruction** |
| Symmetry imposed | D1 | | D5 | |
| Number of particle images | 454,084 | 577,289 | 697,396 | 809,111 |
| Helical twist (°) | 81.1 | | -27.6 | |
| Helical rise (Å) | 19.2 | | 84.2 | |
| Resolution (Å) | 4.04 | 3.26 | 3.98 | 3.31 |
| Map sharpening B factor (Å$^2$) | -230 | -97.4 | -230 | -101.7 |
| FSC threshold | 0.143 | 0.143 | 0.143 | 0.143 |

| **Refinement** | **Narrow helical tube** | | **Wide helical tube** | |
|---|---|---|---|---|
| | **Helical reconstruction** | **Local reconstruction** | **Helical reconstruction** | **Local reconstruction** |
| EMD accession code | EMD-47516 | EMD-42831 | EMD-47517 | EMD-42790 |
| PDB accession code | 9E4N | 8UZ2 | 9E4O | 8UXZ |
| Model composition | | | | |
| Non-hydrogen atoms | 17,528 | 17,528 | 17,528 | 17,528 |
| Protein residues | 2,252 | 2,252 | 2,252 | 2,252 |
| Ligand | 10 | 10 | 10 | 10 |
| Ramachandran plot | | | | |
| Favored (%) | 95.03 | 95.21 | 91.02 | 95.83 |
| Allowed (%) | 4.97 | 4.79 | 8.98 | 4.17 |
| Disallowed (%) | 0.00 | 0.00 | 0.00 | 0.00 |
| Validation | | | | |
| MolProbity score | 1.85 | 1.47 | 2.06 | 1.45 |
| Clashscore | 8.90 | 3.11 | 9.41 | 3.45 |
| Rotamer outliers (%) | 0.00 | 0.00 | 0.00 | 0.00 |
| R.m.s. deviations | | | | |
| Bond lengths (Å) | 0.003 | 0.004 | 0.004 | 0.003 |
| Bond angles (°) | 0.634 | 1.022 | 0.677 | 0.539 |
| Cβ outliers (%) | 0.00 | 0.00 | 0.00 | 0.00 |

[a] Approximate range of underfocus.
